# Cullin-RING ligase BioE3 reveals molecular-glue-induced neosubstrates and rewiring of the endogenous Cereblon ubiquitome

**DOI:** 10.1101/2025.01.07.631759

**Authors:** Laura Merino-Cacho, Orhi Barroso-Gomila, Mónica Pozo-Rodríguez, Veronica Muratore, Claudia Guinea-Pérez, Álvaro Serrano, Coralia Pérez, Sandra Cano-López, Ainhoa Urcullu, Mikel Azkargorta, Ibon Iloro, Carles Galdeano, Jordi Juárez-Jiménez, Ugo Mayor, Felix Elortza, Rosa Barrio, James D. Sutherland

**Author notes:** Biobizkaia Health Research Institute, 48903 Barakaldo, Spain (Orhi Barroso-Gomila). Telethon Institute of Genetics and Medicine (TIGEM), Pozzuoli, Italy; Department of Clinical Medicine and Surgery, Federico II University, Naples, Italy.

## Abstract

**Background:** The specificity of the ubiquitination process is mediated by the E3 ligases. Discriminating genuine substrates of E3s from mere interacting proteins is one of the major challenges in the field. We previously developed BioE3, a biotin-based approach that uses BirA-E3 fusions together with ubiquitin fused to a low-affinity AviTag to obtain a site-specific and proximity-dependent biotinylation of the substrates. We proved the suitability of BioE3 to identify targets of RING and HECT-type E3 ligases.

**Methods:** BioE3 experiments were performed in HEK293FT and U2OS stable cell lines expressing TRIPZ-bio^GEF^Ub transiently transfected with BirA-cereblon (CRBN). Cells were seeded using biotin-free media, adding a short-biotin pulse. We evaluated the applicability of the BioE3 system to CRBN and molecular glues by western blot and confocal microscopy, blocking the proteasome with bortezomib, inhibiting NEDDylation with MLN4924 and treating the cells with pomalidomide. For the identification of endogenous substrates and neosubstrates we analyzed the eluates of streptavidin pull-downs of BioE3 experiments by LC-MS/MS. Analysis of targets which ubiquitination changes significantly upon treatment was done using two-sided Student’s t-test. Orthogonal validations were performed by histidine pull-down, GFP-trap and computational modelling.

**Results:** Here we demonstrate that BioE3 is suitable for the multi-protein complex Cullin-RING E3s ligases (CRLs), the most utilized by targeted protein degradation strategies. Choosing CRBN as proof of concept, one of the substrate receptors of CRL4 E3 ligase, we identified both endogenous substrates and novel neosubstrates upon pomalidomide treatment, including CSDE1 which contains a G-loop motif potentially involved in the binding to CRBN in presence of pomalidomide. Importantly, we observed a major rearrangement of the endogenous ubiquitination landscape upon treatment with this molecular glue.

**Conclusions:** The ability of BioE3 to detect and compare both substrates and neosubstrates, as well as how substrates change in response to treatments, will facilitate both target and off-target identifications and offer a broader characterization and validation of targeted protein degradation degraders, like molecular glues and PROTACs.

## Background

Protein ubiquitination is a post-translational modification involved in almost all cellular processes that plays a crucial role in the regulation of protein homeostasis (1). Ubiquitin (Ub) is covalently attached to the target protein in an highly regulated enzymatic cascade that involves two activating E1s, around 40 conjugating E2s and more than 600 E3 ligases (2). Furthermore, ubiquitination can be reversed by deubiquitinating enzymes (DUBs) (3). Depending on their mechanism of Ub transfer to the substrate protein, E3s are classified in three main families: RING (Really Interesting New Gene; around 600 members), HECT (Homology to E6AP C Terminus, around 30 members) and RBR (RING-Between-RING, around 15 members) (4, 5).

Cullin-RING E3 ligases (CRLs) are the most abundant class of RING E3s and are composed by multiple subunits (6). These complexes contain a cullin scaffold, a RING finger protein (RBX1 or RBX2) that binds the E2, and distinct sets of adaptors and substrate receptors that specifically recruit target proteins. CRL activity requires cullin NEDDylation and is downregulated by deNEDDylation mediated by the COP9 signalosome. CRLs are presently the most used subfamily for targeted protein degradation (TPD), a strategy that uses chemicals to recruit undesired proteins-of-interest to an E3 ligase for ubiquitination and degradation. CRLs based on cereblon (CRBN) and Von Hippel-Lindau tumor suppressor (VHL) are currently the most commonly used in TPD. Specifically, CRBN is the substrate receptor of the CUL4–RBX1–DDB1–CRBN (CRL4^CRBN^) E3 complex. CRBN was identified as the target of immunomodulatory imide drugs (IMiDs) (7), which include thalidomide, pomalidomide and lenalidomide. These compounds and their derivatives have been the basis for many described protein degradation drugs, like monovalent molecular glues (MGs) and bivalent proteolysis targeting chimeras (PROTACs) (8, 9). IMiDs bind to CRBN and alter its substrate specificity, acting as MGs and leading to ubiquitination of non-native substrates (neosubstrates). Ubiquitination by a particular E3 or on particular substrates can occur in diverse ways (on a single site, multiple sites, and with extended ubiquitin-chains of different linkages and topologies). This influences substrate fate and often (but not always) leads to protein degradation. Therefore, to map the profile of endogenous ubiquitinated substrates and of neosubstrates upon cellular treatment with molecular glues and PROTACs should be a crucial step in their validation.

Understanding substrate recognition by particular E3 ligases is a relevant area of research in the Ub field, especially in the light of new developments in TPD (10). Assays that aid in the discovery and/or characterization of substrate specificity of E3 ligases will be an important addition to the chemical biology and drug discovery toolbox. Different strategies to identify targets of E3 ligases have been developed. Proximity proteomics, which has been applied to members of the multi-subunit RING SCF (Skp, Cullin, F-box) complex and others, may identify interactors, some of which may be targets (11–13). Other strategies involve the direct fusion of E3 ligases to Ub-like proteins (UbLs) (UBAIT, TULIP, and SATT) or Ub-binding domains (14–17). Overexpression of an E3 in combination with tagging of Ub has also been used to identify candidate E3 substrates (18, 19), but some of these substrates might not be direct. To complement these approaches, we recently developed BioE3 (20), a biotin-based strategy based on two elements: (1) the fusion of the BirA enzyme, a biotin ligase that labels specifically a biotin acceptor peptide (AviTag), to the E3 ligase of interest; and (2) a UbL fused to an AviTag with lower affinity for BirA (bio^GEF^) (21). The use of bio^GEF^UbLs allows a site-specific and proximity-dependent biotinylation that leads to the specific labeling of the ubiquitinated substrates. Those can be captured by streptavidin pull-down and identified using liquid chromatography-mass spectrometry (LC-MS/MS) proteomics. BioE3 was applied to the RING non-associated to cullins (RNF4, MIB1, MARCH5 and RNF214) and HECT (NEDD4) E3 ligases. Similar methods (E-STUB and Ub-POD) have been described that support this bioUb-based approach to identify targets (22, 23).

Here we demonstrate how BioE3 can be used for the identification of both CRBN endogenous substrates and neosubstrates in presence of iMiDs. By fusing BirA to the N-terminus of CRBN, we show specific biotinylation in HEK293FT- and U2OS-TRIPZ-bio^GEF^Ub cells. We validated Spalt-like 4 (SALL4) as a neosubstrate upon pomalidomide treatment, thus confirming the capacity of CRBN BioE3 to identify neosubstrates. Our proteomic study identified known and novel endogenous substrates of CRBN, and potential pomalidomide-induced neosubstrates, including CSDE1 (Cold Shock Domain Containing E1), with orthogonal validation and computational modelling to explore binding sites. Importantly, we discovered global differences in the ubiquitination of endogenous substrates upon pomalidomide treatment. By revealing changes in both endogenous substrates and neosubstrates of particular E3-drug combinations, we anticipate that BioE3 will be a very useful tool in the future development of TPD.

## Methods

### Cell Culture

U2OS (ATCC HTB-96) and HEK293FT (Invitrogen) were cultured at 37°C and 5% CO2 in Dulbecco’s Modified Eagle Medium (DMEM) supplemented with 10% fetal bovine serum (FBS, Biowest) and 1% penicillin/streptomycin (Gibco). HEK293FT cells were used for Western blot and mass spectrometry experiments, whereas U2OS cells were used for confocal microscopy. For all BioE3 experiments, cells were pre-cultured for 24 hours in biotin-free media supplemented with 10% dialyzed FBS (3.5 kDa MWCO; 150 mM NaCl; filter-sterilized) prior to transfections to control biotin labelling timings. Cultured cells were maintained for maximum 20 passages and tested negative for mycoplasma.

### Cloning

Plasmids were generated by standard cloning or Gibson Assembly (NEBuilder HiFi Assembly, NEB). XL10-Gold bacteria (Agilent) were used. Depending on the construction, we used plasmid backbones derived from TRIPZ (Open Biosystems/Horizon) or Lenti-Cas9-blast (Addgene #52962, kindly provided by F. Zhang). TRIPZ-bio^GEF^Ub and TRIPZ-bio^GEF^Ubnc were previously described (Addgene #208045, 208044) (20). CRBN ORF was amplified from hTERT-RPE1 cell cDNA by high-fidelity PCR (Platinum SuperFi DNA Polymerase; Invitrogen #12351010) and was inserted into the *Eco*R1-*Not*1 sites of Lenti-EFS-BirAopt-GSQ-RBXN-P2A-blast (Addgene #208048) (20). CRBN mutation described in the text was introduced by 2-fragment overlap PCR and Gibson assembly or using primers: CRBN.W386A.qc.for (agctggtttcctgggtatgccGCTactgttgcccagtgtaagatc) and CRBN.W386A.qc.rev (gatcttacactgggcaacagtAGCggcatacccaggaaaccagct). Constructions were validated by Sanger sequencing. Further details will be available upon request.

### Lentiviral transduction

Packaging of lentiviral expression constructs was done in HEK293FT cells by transfecting psPAX2 and pMD2.G (kindly provided by D. Trono; Addgene #12260, #12259) and pTAT (kindly provided by P. Fortes; for TRIPZ-based vectors) using calcium phosphate. After 12-18 hours transfection media were removed and replaced with fresh media. Lentiviral supernatants were collected twice (24 hours each), pooled, filtered (0.45 µm), supplemented with sterile 8.5% PEG6000, 0.3 M NaCl, and incubated for 12-18 hours at 4°C. Lentiviral particles were concentrated by centrifugation (1, 500 x g, 45 minutes, 4°C). HEK293FT and U2OS cells were transduced with non-concentrated or 5x concentrated virus, respectively. Drug selection was performed with 1 µg/ml puromycin (ChemCruz).

### Transfections and drug treatments

HEK293FT and U2OS cells were transfected using calcium phosphate or Lipofectamine 3000 (Thermo Fisher), respectively. TRIPZ cell lines stably transduced were induced with DOX (doxycycline hyclate 1 µg/ml; 24 hours; Sigma-Aldrich) prior to biotin treatment (50 µM; 2 hours; Sigma-Aldrich). BTZ (200 nM; MedChemExpress), MLN4924 (1 µM, MedChemExpress) and pomalidomide (10 µM, MedChemExpress) treatments were performed without biotin prior to cell lysis or immunostaining at the indicated time-points.

### Western blot analysis

To remove excess biotin, we washed cells with 1x PBS and then we lysed them in highly stringent washing buffer (WB) 5 (WB5: 8M urea, 1% SDS in 1x PBS) containing 1x protease inhibitor cocktail (Roche) and 50 µM N-Ethylmaleimide (NEM, Alfa Aesar). Samples were sonicated and centrifuged (16, 000 x g, 30 minutes at room temperature, RT). Protein concentration was determined by the BCA Protein Assay (Pierce) following manufacturer’s instructions. For SDS-PAGE, 20 µg of protein were loaded and transferred to nitrocellulose membranes. PBT (1x PBS, 0.1% Tween-20) was used for blocking, except for anti-biotin blots, where casein-based blocking solution (Sigma) was used. Primary antibodies were incubated for 2 hours at RT or overnight at 4°C and secondary antibodies for 45 minutes at RT. Antibodies were used as follows: anti-biotin-HRP (1/1, 000, Cell Signaling Technology Cat#7075S); anti-BirA (1/1, 000, SinoBiological Cat#11582-T16); anti-AviTag (1/1, 000, GenScript Cat#A00674); anti-NEDD8 (1/1, 000, Abcam Cat# ab81264); anti-GAPDH (1/5, 000, Proteintech Cat# 60004-1-Ig); anti-CSDE1 (1/1, 000, Proteintech Cat# 13319-1-AP); anti-GFP (1/1, 000, Roche Cat#11814460001); anti-GFP (1/2, 000, rabbit polyclonal serum; generated in-house against recombinant GFP protein); anti-HA-tag (1/2, 000, Cell Signaling Technology Cat#3724) anti-Mouse-HRP (1/5, 000, Jackson ImmunoResearch Cat#115-035-062), anti-Rabbit-HRP (1/5, 000, Jackson ImmunoResearch Cat#111-035-045). We used Super Signal West Femto (ThermoFisher) or Clarity ECL (BioRad) to detect the proteins in an iBright CL1500 (Thermo Fisher). Uncropped blots are provided as Supplementary file 1.

### Immunostaining and confocal microscopy

U2OS cells were seeded on 11 mm coverslips (20, 000 cells per well; 24 well plate). After washing the cells with 1x PBS they were fixed with 4% PFA supplemented with 0.1% Triton X-100 in 1x PBS for 20 minutes at RT. Then, coverslips were washed 3 times with 1x PBS and blocked in blocking buffer (2% fetal calf serum, 1% BSA in 1x PBS) for 30 minutes at RT. Primary antibodies were incubated for 1 hour at 37°C and cells were washed 3 times with 1x PBS. Primary antibodies were used as follow: anti-BirA (1/200, Novus Biologicals Cat#NBP2-59939); anti-NEDD8 (1/100, Abcam Cat# ab81264). Secondary antibodies and fluorescent streptavidin were incubated for 30 minutes at 37°C. After that, nuclei were stained with DAPI (300 ng/ml in 1x PBS, Sigma Aldrich) for 10 minutes. Secondary antibodies (ThermoFisher) were all used at 1/200: anti-Mouse Alexa Fluor 488 (Cat#A-11029), anti-Rabbit Alexa Fluor 647 (Cat#A-21244), anti-Mouse Alexa Fluor 647 (Cat#A-31571). Streptavidin Alexa Fluor 594 (1/200, Cat#016-290-084, Jackson ImmunoResearch) was also used. Images were taken with a confocal microscope (Leica SP8 Lightning) using 63x Plan ApoChromat NA1.4 objective.

### Pull-down of biotinylated proteins

The lysates cleared in WB5 were normlized to the same protein concentration and incubated overnight at RT with equilibrated NeutrAvidin-agarose beads (ThermoFisher) at a ratio of 1/50 (v_beads_/ v_lysate_). The high affinity between biotin and streptavidin allows stringent series of washes, as follows (v_WB_/2v_lysate_): 2x WB1 (8 M urea, 0.25% SDS); 3x WB2 (6 M Guanidine-HCl); 1x WB3 (6.4 M urea, 1 M NaCl, 0.2% SDS); 3x WB4 (4 M urea, 1 M NaCl, 10% isopropanol, 10% ethanol and 0.2% SDS); 1x WB1; 1x WB5; and 3x WB6 (2% SDS; WB1-6 prepared in 1x PBS). Biotinylated proteins were eluted from the beads using 1 volume of Elution Buffer (4x Laemmli buffer, 100 mM DTT; 80 µl for LC-MS/MS experiments) by heating at 99°C for 5 minutes twice, followed by vortexing. Beads were separated using clarifying filters (2, 000 x g, 2 minutes; Vivaclear Mini, Sartorius).

### Liquid Chromatography Mass Spectrometry (LC-MS/MS)

Pull-down experiments for mass-spectrometry were performed independently in triplicates. For each replicate, four confluent 15 cm dishes (8 × 10^7^ cells, 2 ml of lysis per plate; 8 ml total) were analyzed by LC-MS/MS. Samples eluted from the NeutrAvidin beads were separated in SDS-PAGE and stained with Sypro Ruby (Invitrogen) following manufacturer’s instructions. Gel lanes were carefully cut to ensure consistency and reproducibility. Slices were subsequently washed in milli-Q water. Reduction and alkylation were performed (10 mM DTT in 50 mM ammonium bicarbonate, 56°C, 20 minutes, followed by 50 mM chloroacetamide in 50 mM ammonium bicarbonate, 20 minutes, protected from light). Gel pieces were dried and incubated with trypsin (12.5 µg/ml in 50 mM ammonium bicarbonate, 20 minutes, ice-cold). After rehydration, the trypsin supernatant was discarded. After hydration with 50 mM ammonium bicarbonate, gel pieces were incubated at 37°C overnight. Following digestion, 0.1% TFA was used to clean acidic peptides, which were dried in a RVC2 25 SpeedVac concentrator (Christ). Peptides were resuspended in 10 µl 0.1% formic acid (FA) and sonicated for 5 minutes prior to analysis.

Samples were analyzed using a timsTOF Pro mass spectrometer (trapped ion mobility spectrometry/quadrupole time of flight hybrid, Bruker Daltonics) coupled online to a EVOSEP ONE (Evosep), which uses parallel accumulation–serial fragmentation (PASEF), at the proteomics platform of CIC bioGUNE. Sample (200 ng) was directly loaded in a 15 cm performance column (Evosep) applying a 30 samples per day method and data dependent acquisition mode.

### Mass Spectrometry data analysis

DIA data was processed with DIA-NN software for protein identification and quantification using default parameters. Searches were carried out against a database consisting of *Homo sapiens* protein entries from Uniprot in library-free mode. We considered carbamidomethylation of cysteines and oxidation of methionines as fixed and variable modifications, respectively. Match between runs was applied and precursor FDR was set at 1%. Data was processed and analyzed by Perseus (version 1.6.15) (24). Proteins identified by at least 2 peptides and present in at least 2 out of 3 replicates in at least one group were included in the analysis. Statistical significance was assessed using a two-sided Student’s t-test. Data were loaded into GraphPad Prism 10 version 10.0.2 to build the corresponding volcano-plots.

Network analysis was conducted using STRING version 1.4.2 in Cytoscape version 3.9.1, applying a high-confidence interaction score of 0.7 (25, 26). Size, transparency and width of the edges were continuously mapped to the Log2 fold change. The Molecular COmplex DEtection (MCODE) plug-in version 1.5.1 was used to identify highly connected subclusters of proteins (degree cutoff of 2; Cluster finding: Haircut; Node score cutoff of 0.2; K-Core of 2; Max. Depth of 100) (27). Gene ontology analysis was performed using g:Profiler web server version e108_eg55_p17_0254fbf and REVIGO (28, 29). Venn diagrams were drawn using InteractiVenn web tool (30).

### Histidine pull-down

For orthogonal validations, HEK293FT cells were co-transfected with the indicated constructs and pcDNA3-His6-Ub (gift from M. Rodriguez, CRNS-LCC, Toulouse), lysed in lysis buffer (8 M urea, 0.1 M Na2HPO4/NaH2PO4 pH 8.0, 0.01 M Tris-HCl pH 8.0, 20 mM imidazole pH 8.0, 5 mM β-mercaptoethanol, and 0.1% Triton X-100), supplemented with 1x protease inhibitor cocktail (Roche) and 50 µM PR619 DUB inhibitor (Merck). Samples were then sonicated and cleared by centrifugation (25, 000 x g, 30 minutes at RT). Cleared lysates were adjusted to the same protein concentration before incubating them with 1/50 (vol_beads_/vol_lysate_) equilibrated Ni-NTA agarose beads (Invitrogen) overnight at RT. Beads were then washed three times using WBB (8 M urea, 0.1 M Na_2_HPO_4_/NaH_2_PO_4_ pH 8.0, 0.01 M Tris-HCl pH 8.0, 20 mM imidazole pH 8.0, 2.5 mM β-mercaptoethanol, and 0.1% Triton X-100), and two times using WBC (8 M urea, 0.1 M Na_2_HPO_4_/NaH_2_PO_4_ pH 6.3, 0.01 M Tris-HCl pH 6.3, 10 mM imidazole pH 7.0, 2.5 mM β-mercaptoethanol, and 0.1% Triton X-100). Proteins were eluted with 1 vol_beads_ of Elution Buffer (4 M urea, 50 mM NaH_2_PO_4_/Na_2_HPO_4_, 5 mM Tris/HCl pH 8, 500 mM, imidazole pH 7.0, 1.25 mM β-mercaptoethanol, and 0.05% Triton X-100).

### GFP-trap pull-down

All steps were performed at 4°C. HEK293FT cells were collected 48 hours after transfection, washed 3 times with 1x PBS and lysed in RIPA lysis buffer (50 mM Tris-HCl pH 8, 150 mM NaCl, 1% IGEPAL CA-630, 0.5% sodium deoxycholate, and 0.1% SDS) supplemented with 1x protease inhibitor cocktail (Roche), BTZ (MedChemExpress) and 50 µM PR619 (Merck). Lysates were kept on ice for 30 minutes and cleared by centrifugation (25, 000 x g, 30 minutes at 4°C). Cleared lysates were incubated with 15 µl of equilibrated GFP Selector beads (Nanotag Biotechnologies) overnight at 4°C in a rotating wheel. Beads were washed 4 times with RIPA lysis buffer, twice with RIPA/4 M urea, and once more with RIPA. The samples were eluted in 2x Laemmli buffer by boiling for 5 minutes at 95°C.

### Molecular Modelling

The G-loop regions of CSDE1 were identified by comparing the position of alpha carbon atoms between the CSDE1 AlphaFold model AF-O75534-F1 and the crystallographic structure of SALL4 with PDB id: 7BQU. The structure for the first CSDE1 domain was obtained from the AlphaFold model AF-O75534-F1 and the second domain was extracted from the NMR model with PDB id: 2YTV. The ternary complexes were modeled by superimposing each of the G-loop sequences of CSDE1 to the structure of CK1α in complex with CRBN (PDB id 5FQD) as reference and the pomalidomide-binding mode in the thalidomide-binding domain was taken from the crystallographic structure with PDB id: 6H0F. The AMBERff14SB and GAFF2 force fields were used to assign atom types for the proteins and pomalidomide, respectively. The partial charges for pomalidomide were derived using the RESP protocol at the HF/6-31G(d) level of theory, calculated using Gaussian16 (31). The Zn2+ cation coordinated with CRBN was modeled with the bound model, using the ZAFF parameters (32). The systems were solvated on a truncated octahedral box of TIP3P water molecules, and neutralized with salt counterions. Following the protocol we have recently reported (33), each system was minimized, heated to 298K, and equilibrated to 1 bar. Each H-bond in the ternary complex interface was evaluated using 100 independent steered molecular dynamics trajectories. Starting positions and velocities for each system were sampled from independent classical MD 10 ns trajectories, using a flat-bottom restraint to keep the H-bonds between 2.5 and 3.5 Å. Then the H-bonds were brought to a 2.5 Å distance to start a constant speed steering at a speed of 0.5 Å/ns, using the stiff spring approximation. All simulations were performed with the CUDA accelerated version of PMEMD from the Amber23 package (34). The potential mean force (PMF) of each H-bond was computed using the Jarzynski equality on the resulting work profiles (35). The error estimations for the PMF profiles were obtained by bootstrapping 25 times with 25 replica subsamples.

## Results

### Applying BioE3 to study Cullin-RING E3 ligases (CRL)

We previously used BioE3 for detecting specific targets of RING and HECT type E3 ligases (20), but the potential of BioE3 to detect targets of E3s multi-protein complexes like the CRLs was not tested. First, we considered the localization of the BirA enzyme, which could produce steric problems. We performed BioE3 experiments with BirA fused to the N-or C-terminus of CRBN and we observed similar pattern of biotinylated proteins (Fig. S2a; Supplementary file 1). Based on previous reports (36, 37), we considered the localization of the BirA fusion to the N-terminus of CRBN (Fig. 1a, b) and transiently transfected it into stable cell lines expressing the bio^GEF^Ub in a doxycycline (DOX)-dependent manner (TRIPZ-bio^GEF^Ub). The low-affinity bio^GEF^ AviTag enables site-specific and proximity-dependent biotinylation (21). In this way, after DOX induction for 24 hours and controlled biotin pulses, substrates modified by the CRL4^BirA-CRBN^ and labelled with biotinylated bio^GEF^Ub can be purified using streptavidin pull-down for identification by LC-MS/MS, or imaged by immunofluorescence and confocal microscopy (Fig. 1a, b).

**Fig. 1.**
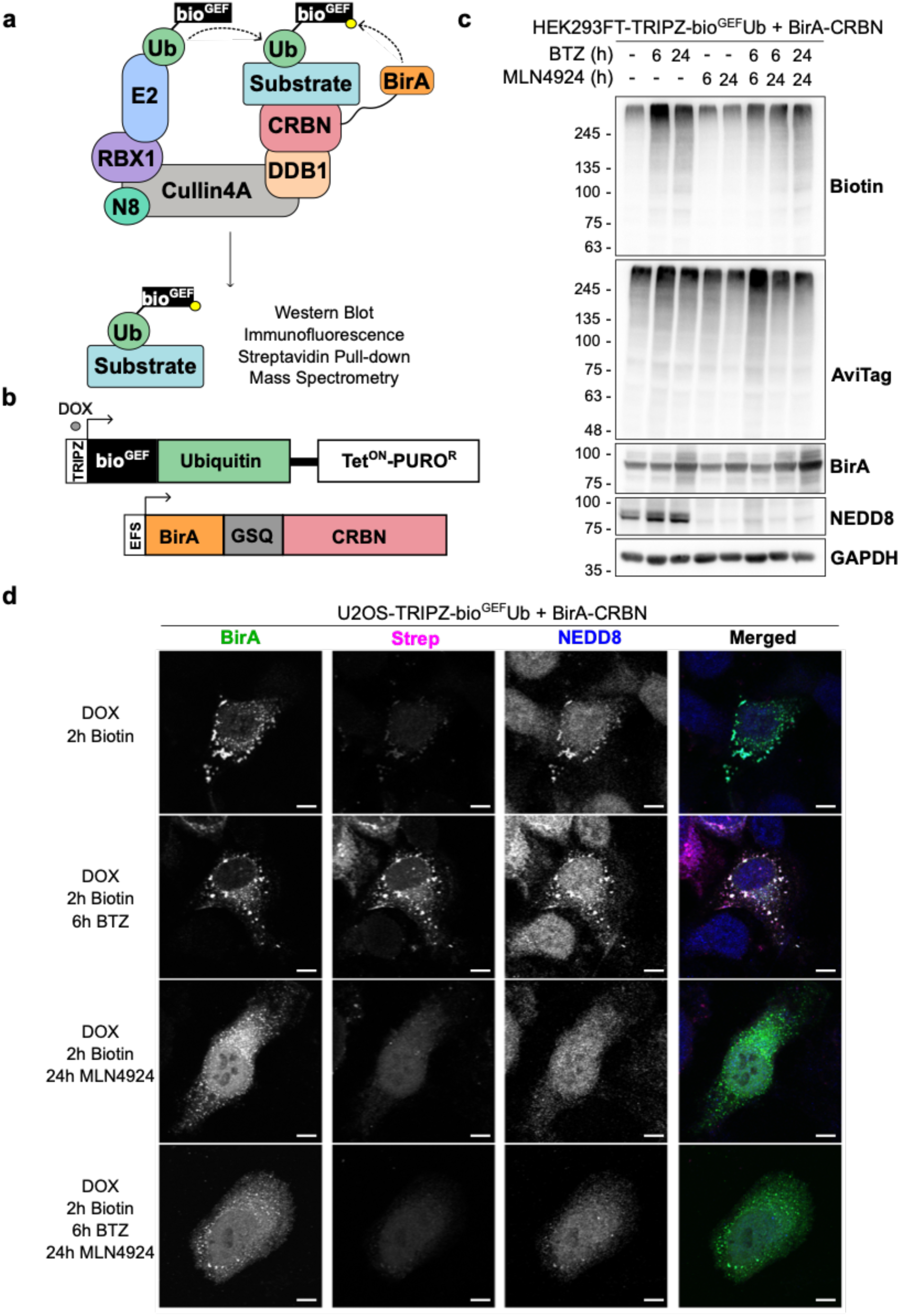
BioE3 labels CRL-dependent ubiquitinated substrates of CRBN. **a, b)** Schematic representation of the BioE3 strategy adapted to the substrate receptor CRBN **(a)** and the constructs used in this work **(b)**. TRIPZ, all-in-one inducible lentiviral vector; bio^GEF^, low affinity AviTag (see text); DOX, doxycycline; Tet^ON^, tetracycline inducible promoter; PURO^R^, puromycin resistant cassette; EFS, elongation factor 1α short promoter. **c)** Western blot of BioE3 experiment performed on HEK293FT stable cell line expressing TRIPZ-bio^GEF^Ub and transfected with EFS-BirA-CRBN. Indicated samples were treated with 100 nM bortezomib (BTZ) for 24 hours, 200 nM BTZ for 6 hours and/or 1 µM MLN4924 for 6 or 24 hours. Molecular weight markers are shown to the left of the blots in kDa, antibodies used are indicated to the right. AviTag antibodies highlight all the ubiquitinated proteins, while biotin shows those ubiquitinated by CRBN. **d)** Confocal microscopy images of BioE3 experiment performed on U2OS stable cell line expressing TRIPZ-bio^GEF^Ub transfected with BirA-CRBN. Indicated samples were treated with 200 nM BTZ for 6 hours and/or 1 µM MLN4924 for 24 hours. Biotinylated material is stained with fluorescent streptavidin (Strep, magenta) and BirA (green) and NEDD8 (blue) with specific antibodies. Scale bar: 8 µm. All BioE3 experiments were performed by pre-incubating the cells in dialyzed FBS-containing media prior to transfections, doxycycline (DOX) induction at 1 µg/ml for 24 hours and biotin supplementation at 50 µM for 2 hours.

To test the specificity of the system, we performed BioE3 using BirA-CRBN and HEK293FT-TRIPZ-bio^GEF^Ub cells, inducing the expression of bio^GEF^Ub with DOX and performing a 2-hour biotin pulse. We observed biotinylated proteins, indicating the activity of BirA-CRBN fusion (Fig. 1c; Supplementary file 1), similarly to what was previously reported by Huang and collaborators for CRBN-BirA (22). Additionally, when proteasome activity was blocked with bortezomib (BTZ) we observed a further accumulation of biotinylated proteins that was reversed upon inhibition of NEDDylation with MLN4924 (Fig. 1c).

We then confirmed co-localization of the BirA-CRBN fusion protein with NEDD8 in the cytoplasm by confocal microscopy in U2OS-TRIPZ-bio^GEF^Ub cells (Fig. 1d), which suggest that the fusion was correctly incorporated into the CRL complex. Upon proteasomal inhibition we observed an increase in biotinylated proteins (Fig. 1d, Strep panel) that co-localized with the BirA enzyme, indicating specific biotinylation. MLN4924 treatment reduced the biotin labeling and dispersed the NEDD8 signal. Taken together, these data suggest that the BioE3 system is biotinylating CRBN substrates in a CRL-dependent manner.

Next, we tested wild type (WT) bio^GEF^Ub for its use in CRBN-BioE3. In previous experiments (20), we used a non-cleavable version of Ub bearing the L73P mutation (Ubnc), to prevent the recycling of biotinylated bio^GEF^Ub by suppressing access of DUBs. However, to analyze HECT-type E3s (e.g. NEDD4), we found that it is necessary to use the bio^GEF^Ub, since bio^GEF^Ubnc was not efficiently passed from E2 to E3 (20). We performed BioE3 experiments both in HEK293FT-TRIPZ-bio^GEF^Ub and HEK293FT-TRIPZ-bio^GEF^Ubnc stable cell lines (Fig. S2b) and observed higher abundance of biotinylated proteins when using bio^GEF^Ub. Importantly, as shown in Fig. 1d, biotinylated material was restricted to BirA-CRBN localization, indicating the specificity of labelling in our conditions, so bio^GEF^Ub was used for rest of experiments shown here.

### BioE3 identifies SALL4 as a neosubstrate of CRBN

The transcription factor SALL4 is one of the best-known neosubstrates of CRBN induced by pomalidomide (38–40), so we aimed to identify SALL4 by BioE3 upon treatment with pomalidomide as a proof of concept. As a negative control, we used a CRBN mutant deficient in IMiD-binding (BirA-CRBN^W386A^) (41) that should act as a negative control for the identification of neosubstrates upon pomalidomide treatment. We performed BioE3 experiments in HEK293FT- and U2OS-TRIPZ-bio^GEF^Ub cell lines transiently transfected with the WT or mutant versions of BirA-CRBN, blocking the proteasome with BTZ and treating the cells with the IMiD drug pomalidomide (Fig. S2c; Supplementary file 1). By Western blot we noticed a change in the biotinylation pattern when treating the samples with pomalidomide, suggesting that BioE3 is sensitive to molecular glues and can differentiate the endogenous targets from the neosubstrates. BirA-CRBN^W386A^ also produced biotinylation of proteins, but without changing the pattern observed after adding the IMiD. By confocal microscopy, we could validate the suitability of the mutant for BioE3 experiments: interestingly, BirA-CRBN^WT^ displayed a nuclear localization after pomalidomide treatment that was not observed with BirA-CRBN^W386A^, together with an increase in the nuclear biotinylated material (Fig. 2a, Strep panel).

**Fig. 2.**
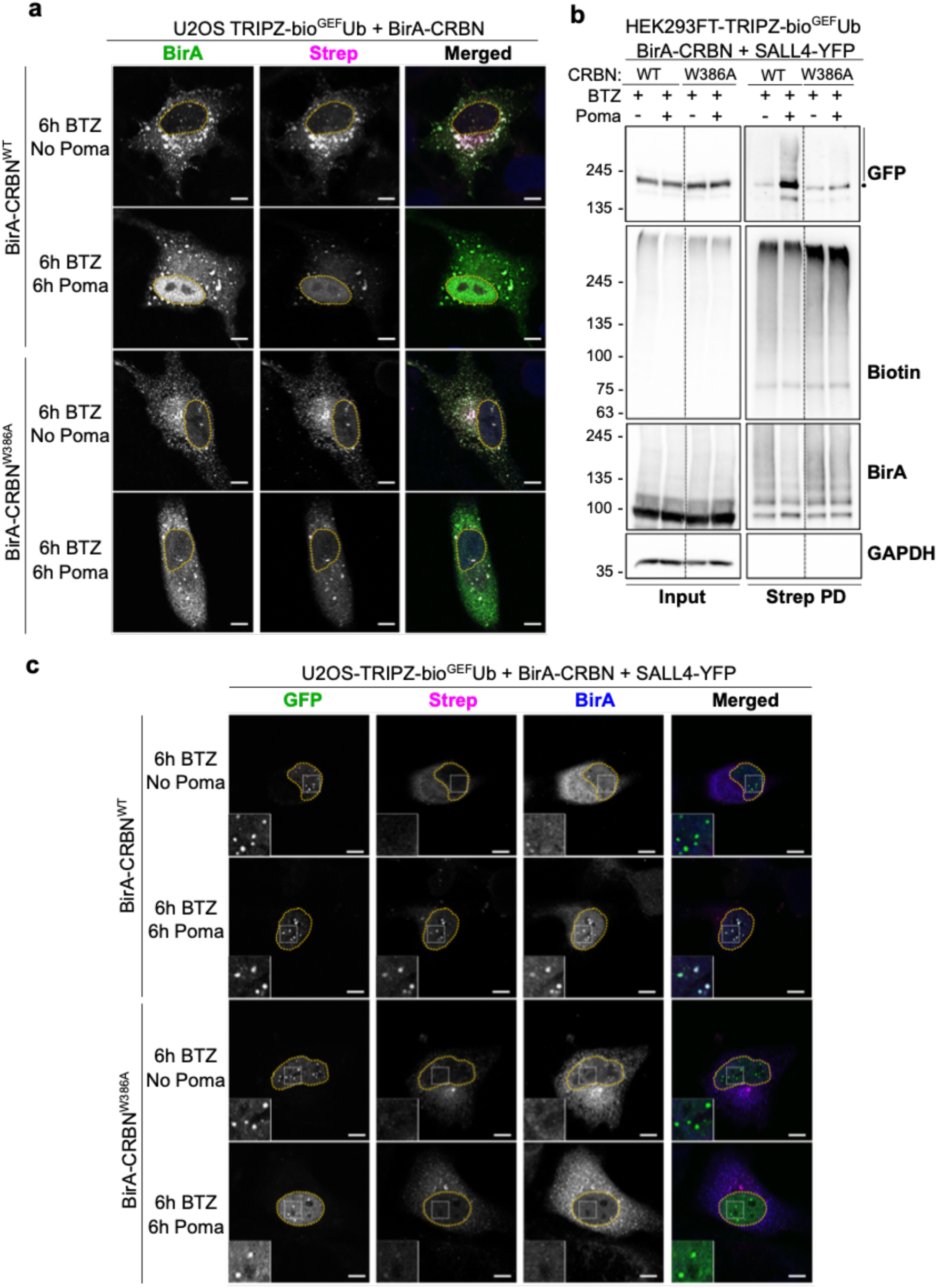
Identification of SALL4 as a neosubstrate of CRBN upon pomalidomide treatment. **a)** Confocal microscopy of BioE3 experiment performed in U2OS stable cell line stably expressing TRIPZ-bio^GEF^Ub. Cells were transfected with EFS-BirA-CRBN^WT^ or EFS-BirA-CRBN^W386A^ (IMiD-binding deficient mutant). Indicated samples were treated with 10 µM pomalidomide (Poma) and/or with 200 nM bortezomib (BTZ) for 6 hours. Biotinylated material is stained with fluorescent streptavidin (Strep, magenta) and BirA (green) with specific antibodies. **b, c)** BioE3 experiment performed in HEK293FT-TRIPZ-bio^GEF^Ub **(b)** or U2OS-TRIPZ-bio^GEF^Ub **(c)** stable cell lines transiently transfected with CMV-SALL4-YFP and EFS-BirA-CRBN^WT^ or EFS-BirA-CRBN^W386A^ and treated with 10 µM pomalidomide (Poma) and/or 200 nM bortezomib (BTZ) for 6 hours. **b)** Western blot validation of SALL4 as a neosubstrate upon pomalidomide treatment. The dot represents the possible monoubiquitinated protein, whereas the bar the polyUb-modified SALL4-YFP. Strep PD: streptavidin pull-down. Molecular weight markers are shown to the left of the blots in kDa, antibodies used are indicated to the right. **c)** Biotinylated material is stained with fluorescent streptavidin (Strep, magenta) and BirA (blue) with a specific antibody. SALL4-YFP can be found in green. Scale bar: 8 µm. Yellow dotted lines indicate the nuclei. Insets show the amplification of the area indicated by a white dotted square in each panel. All BioE3 experiments were performed by pre-incubating the cells in dialyzed FBS-containing media prior to transfections, doxycycline (DOX) induction at 1 µg/ml for 24 hours and biotin supplementation at 50 µM for 2 hours.

Considering that the expression level of SALL4 in HEK293FT cells is low (42), we decided to perform the BioE3 experiment with exogenously expressed SALL4-YFP and treating the cells with BTZ and/or pomalidomide. After isolating the biotinylated proteins by streptavidin pull-down, we observed an enrichment of polyubiquitinated SALL4-YFP upon pomalidomide treatment with the CRBN^WT^ version but not with the IMiD-binding mutant CRBN^W386A^ (Fig. 2b; Supplementary file 1). We confirmed this result using confocal microscopy, observing co-localization of BirA-CRBN^WT^ and biotin signal at nuclear bodies formed by SALL4-YFP only in those cells treated with pomalidomide (Fig. 2c). Furthermore, BirA-CRBN^W386A^ did not co-localize to the SALL4 nuclear bodies, neither biotinylated SALL4-YFP, even after treatment with the IMiD. Altogether, we concluded that BioE3 is sensitive to the responses induced by molecular glues and can be used to discriminate between endogenous substrates and neosubstrates.

### BioE3 identifies endogenous targets of CRBN

Once we confirmed the suitability of BioE3 to biotinylate both putative substrates and neosubstrates after molecular glue treatment, we performed a large-scale experiment in triplicates, treating the cells with BTZ only, BTZ and MLN4924 or BTZ and pomalidomide (Figs. 3, 4; Tables 1-3). We isolated the biotinylated proteins by streptavidin pull-downs and confirmed by Western blot the enrichment of the eluates in biotinylated proteins, when the proteasome was blocked with BTZ (Fig. 3a; Supplementary file 1). By inhibiting NEDDylation the amount of biotinylated proteins was reduced, supporting it as useful negative control. We also observed a reduction in biotinylated proteins after pomalidomide treatment. The eluates were then analyzed by LC-MS/MS to identify specific ubiquitinated targets by CRBN. First, by comparing the samples treated or not with BTZ (BTZ *versus* DMSO) we identified 376 putative targets of CRBN that are proteasome-targeted (Fig. 3b; Supplementary Table 1). Among them, we found glutamine synthetase (GLUL), an endogenous target of the substrate receptor previously described (43).

**Fig. 3.**
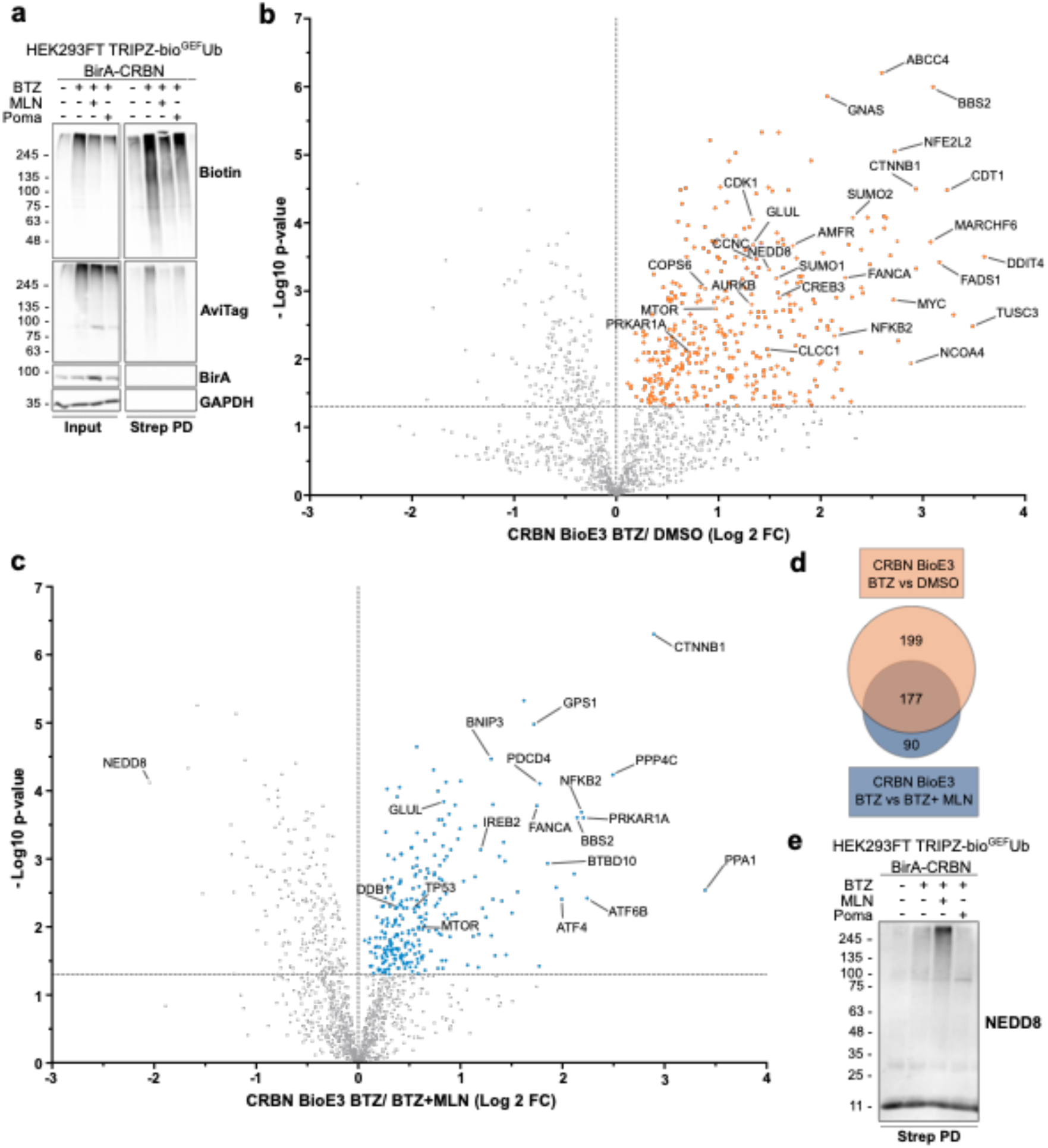
BioE3 identifies endogenous substrates of CRBN. **a)** Western blot of BioE3 experiment performed on HEK293FT stable cell line expressing TRIPZ-bio^GEF^Ub and transfected with EFS-BirA-CRBN. Indicated samples were treated with 200 nM bortezomib (BTZ) for 6 hours, 1 µM MLN4924 for 24 hours and 5 µM pomalidomide (Poma) for 6 hours. **b, c)** Volcano plots of LC-MS/MS analysis comparing streptavidin pull-downs of BioE3 experiments showed in **(a)**. Proteins significantly enriched (Log2 Fold Change (FC) BTZ/DMSO **(b)** or BTZ+MLN4924 **(c)** > 0 and p-value < 0.05) were considered as CRBN targets. Statistical analyses were done using two-sided Student’s t-test. **d)** Venn diagram showing the endogenous targets of CRBN identified by BioE3 in **(b)** and **(c)**. **e)** Western blot of NEDDylated proteins from samples described in **(a)**. Strep PD: streptavidin pull-down. Molecular weight markers are shown to the left of the blot in kDa, antibodies used are indicated to the right. All BioE3 experiments were performed by pre-incubating the cells in dialyzed FBS-containing media prior to transfections, doxycycline (DOX) induction at 1 µg/ml for 24 hours and biotin supplementation at 50 µM for 2 hours.

We also interrogated the effect of inhibiting NEDDylation in the identification of endogenous substrates of CRBN, by comparing BTZ *versus* BTZ and MLN4924 treated cells, and identified 267 putative targets of CRBN that require NEDD8 activation of the CRL complex (Fig. 3c; Supplementary Table 2). Among them, 177 (66%) were also present in the CRBN BioE3 BTZ *versus* DMSO comparison, including GLUL (Fig. 3d). Interestingly, NEDD8 was significantly enriched when inhibiting NAE1 (left quadrant of Fig. 3c), indicating that mixed NEDD8-Ub conjugates were accumulating in that sample. This was also confirmed by Western blot, showing an enrichment in high molecular weight NEDDylated proteins (Fig. 3e; Supplementary file 1). These results are in line with the use of NEDD8 by the Ub machinery when the NEDDylation machinery is blocked (44, 45).

In addition to the aforementioned hits, we identified proteins related to the UbL machinery, including ubiquitination (E3s and DUBs), SUMOylation (SUMO1, SUMO2, PIAS1, PIAS4), NEDDylation (NEDD8, COPS6) and proteasome components. We also found components of the cAMP signaling pathway like MTOR, PRKAR1A and CREB3 (46), components of the Wnt signaling pathway such as CSNK1E and CTNNB1 (47), chloride channels as CLCC1 and CLCN3 (48), BSG for which a ubiquitination-independent, chaperone-like function of CRBN was described (49), and components of CRLs (DDB1) and the COP9 signalosome (COPS6, COPS7A, COPS3) (50).

To determine the functional role of CRBN we performed a STRING network analysis of the potential substrates (Figs. S3 and S5). The network showed a major interconnected core-cluster composed of 71% and 64% of the identified substrates respectively in BTZ *versus* DMSO and BTZ *versus* BTZ and MLN4924. Unsupervised MCODE analysis highlighted sub-clusters related to ribosomes, proteasome, DNA replication, cell cycle and nuclear pore (Figs. S3 and S5). Moreover, the gene ontology analysis showed a significant enrichment in processes related to the Ub Proteasome System (UPS), DNA, cell cycle, apoptosis and autophagy (Figs. S4 and S6; Tables 1 and 2). These results suggest that CRL4^CRBN^ is a highly versatile E3 ligase, involved in different cellular processes, beyond the few already described. In summary, our data support that BioE3 is able to identify endogenous putative targets of CRBN by LC-MS/MS, identifying the cellular processes in which this substrate receptor is implicated.

### BioE3 identifies neosubstrates of CRBN upon pomalidomide treatment by LC-MS/MS

Once we proved the suitability of BioE3 to identify by LC-MS/MS endogenous substrates of CRBN, we analyzed its capacity to identify neosubstrates upon treatment with an IMiD. By analyzing the effect of pomalidomide by LC-MS/MS (BTZ and pomalidomide *versus* BTZ only), we identified 133 neosubstrates (Fig. 4a; Supplementary Table 3). Next, we performed a STRING network analysis defining a major interconnected core-cluster composed of 74% of the putative neosubstrates (Fig. S7). Unsupervised MCODE analysis derived three sub-clusters linked to translation, mRNA processing, chaperones and histones. In the same line, gene ontology analysis highlighted processes related to RNA-binding and translation, actin cytoskeleton, protein folding, chaperones, Ub and apoptosis (Fig. S8; Supplementary Table 3). These results suggest a change in the nature of the neosubstrates in comparison with the endogenous substrates. In agreement with our results, Baek *et al.* identified by affinity purification-MS, novel interactors of CRBN upon IMiD treatment in a recent pre-print (51), the majority of them being non-zinc finger proteins, including RNA-binding proteins. In fact, one of our top hits is CSDE1, an RNA-binding protein implicated in several diseases including several types of cancer (52). CSDE1 was previously reported by Yamanaka *et al*. as part of the pomalidomide-induced CRBN interactome using AirID as a proximity-biotinylation strategy in HEK293FT cells (36). In previous structural characterizations on IMiD-induced ternary complexes (41, 53), a G-loop like structure has been consistently reported as an essential feature for the recruitment of CRBN neosubstrates. Interestingly, CSDE1 contains two G-loop regions (G-loop1: residues 162 to 173; and G-loop2: residues 721 to 732) that closely resemble the CRBN-engaging G-loop of the neosubstrate GSPT1, with a root mean square deviation among the position of alpha carbons < 0.5 Å. Thus, we confirmed by BioE3 the enrichment of ubiquitinated CDSE1 after treating the cells with pomalidomide, but not when the IMiD-binding deficient mutant CRBN^W386A^ was tested (Fig. 4b; Supplementary file 1). Furthermore, we orthogonally validated CSDE1 as a neosubstrate of CRBN by histidine pull-down, observing a correlation in the ubiquitination by the length and concentration of the pomalidomide treatment (Fig. 4c; Supplementary file 1). We also sought to characterize the putative binding of CSDE1 to CRBN by means of molecular modelling. We have previously reported that the strength of H-bonds is an excellent predictor of the stability of ternary complexes between CRBN, IMiDs and the neosubstrate CK1α (33). Thus, we investigated whether the strength of the three hydrogen bonds at the CRBN-CSDE1 interface was modified by the presence of pomalidomide in the IMiD-binding domain of CRBN. We observed that, although two of the three H-bonds at each of the interfaces was largely unaffected (changes on the H-bond strength ∼1.5 kcal/mol) the energy required to break the H-bond between W400 in CRBN and the backbone carbonyl of CSDE1 of T168 (G-loop1) and T727 (G-loop2) was higher by 3.3 +/– 1.0 kcal/mol and 4.6 +/– 1.1 kcal/mol, respectively when pomalidomide was present in the thalidomide-binding domain (Fig. 4d). The increased strength of the H-bond indicates that pomalidomide has a stabilizer effect in the complex between CRBN and CSDE1, although its magnitude is moderate in comparison with other known neosubstrates of CRBN such as CK1α and GSPT1 (41, 53). Altogether, these results suggest that BioE3 can screen for changes in the ubiquitination pattern by CRLs induced by molecular glues, detecting new putative neosubstrates that can be subsequently validated using orthogonal techniques.

**Fig. 4.**
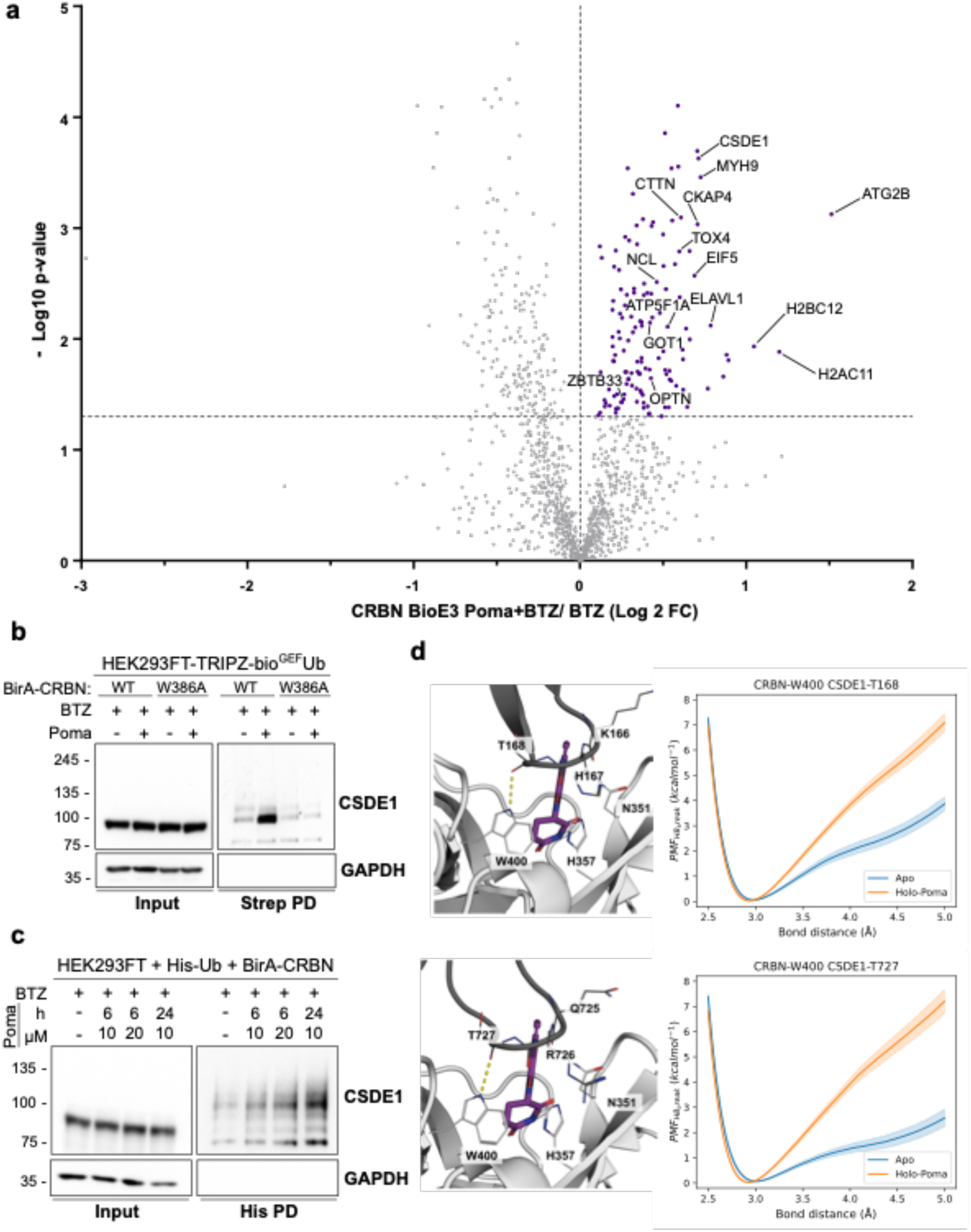
BioE3 identifies neosubstrates of CRBN upon pomalidomide treatment. **a)** Volcano plot of LC-MS/MS analysis comparing streptavidin pull-downs of BioE3 experiments showed in Fig. 3. Proteins significantly enriched (Log2 Fold Change (FC) Poma+BTZ/BTZ > 0 and p-value < 0.05) were considered as CRBN neosubstrates. Statistical analysis was done using two-sided Student’s t-test. **b)** Western blot of a BioE3 experiment performed in HEK293FT-TRIPZ-bio^GEF^Ub stable cell line transiently transfected with EFS-BirA-CRBN^WT^ or EFS-BirA-CRBN^W386A^ and treated with 10 µM pomalidomide (Poma) and/or 200 nM bortezomib (BTZ) for 6 hours. We validated endogenous CSDE1 as a neosubstrate of CRBN upon pomalidomide treatment. All BioE3 experiments were performed by pre-incubating the cells in dialyzed FBS-containing media prior to transfections, doxycycline (DOX) induction at 1 µg/ml for 24 hours and biotin supplementation at 50 µM for 2 hours. **c)** Histidine pull-down confirming CSDE1 as a neosubstrate of CRBN. HEK293FT cells were transiently transfected with BirA-CRBN and His-Ub and treated with 200 nM BTZ for 6 hours and with pomalidomide at the indicated concentrations and time points. Molecular weight markers are shown to the left of the blots in kDa, antibodies used are indicated to the right. **d)** Comparison between the potential mean force (PMF) profiles for the interaction of W400 of CRBN with the backbone carbonyl group of T168 (top panel) and T727 (bottom panel) of CSDE1 in the presence of Poma (orange line) and without ligand in the thalidomide-binding domain (blue line). The solid lines represent the PMF calculated using the Jarzynski equality for 100 independent replicas and the shaded areas represent the standard deviation obtained by bootstrapping.

### Ubiquitination of the endogenous substrates of CRBN changes upon pomalidomide treatment

The effect of IMiD treatment in the ubiquitination pattern of the endogenous substrates remains unknown. Therefore, we studied the ubiquitination of the endogenous substrates that increased, decreased or did not change in the BioE3 analysis upon pomalidomide treatment. Our results showed that most of the endogenous substrates were less ubiquitinated by CRBN, suggesting a major rewiring for CRBN specificity upon molecular glue treatment (Fig. 5a).

**Fig. 5.**
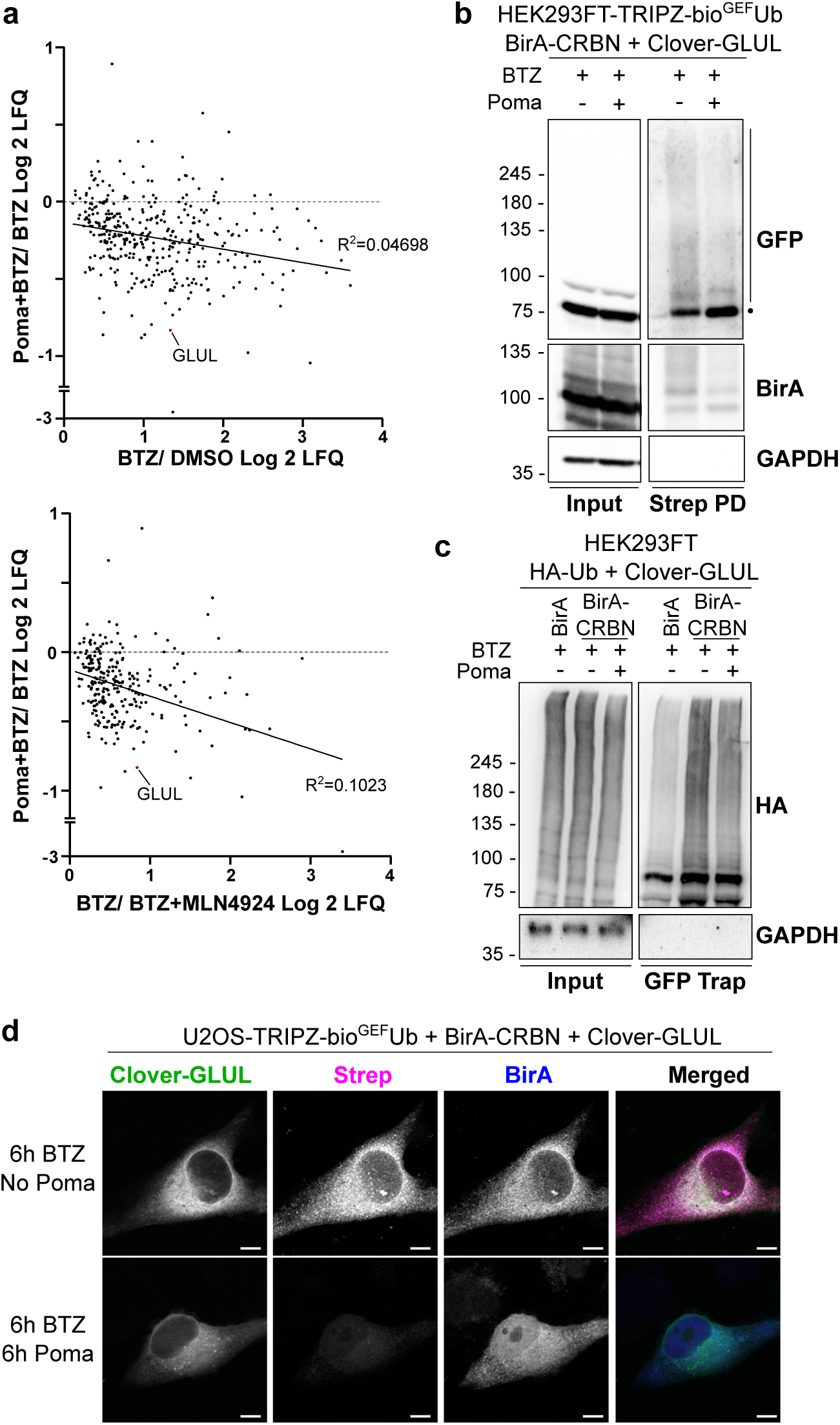
The ubiquitination of endogenous substrates of CRBN decreases upon pomalidomide treatment. **a)** Fold change of the endogenous substrates of CRBN defined in Fig. 3 after treating the cells with pomalidomide. **b, c)** BioE3 experiment performed in HEK293FT-TRIPZ-bio^GEF^Ub **(b)** or U2OS-TRIPZ-bio^GEF^Ub **(c)** stable cell lines transiently transfected with EFS-Clover-GLUL and EFS-BirA-CRBN and treated with 10 µM pomalidomide (Poma) and/or 200 nM bortezomib (BTZ) for 6 hours. **b)** Western blot validation by BioE3 of GLUL as a substrate of CRBN. The dot represents the unmodified protein, whereas the bar the Ub modified. Strep PD: streptavidin pull-down. Molecular weight markers are shown to the left of the blots in kDa, antibodies used are indicated to the right. **c)** GFP-trap pull-down confirming the reduction in the ubiquitination of CRBN upon pomalidomide treatment. HEK293FT cells were transiently transfected with BirA-CRBN, Clover-GLUL and HA-Ub and treated with 200 nM BTZ and with 10 µM pomalidomide for 6 hours as indicated. **d)** Biotinylated material is stained with fluorescent streptavidin (Strep, magenta) and BirA (blue) with a specific antibody. Clover-GLUL can be found in green. Scale bar: 8 µm. All BioE3 experiments were performed by pre-incubating the cells in dialyzed FBS-containing media prior to transfections, doxycycline (DOX) induction at 1 µg/ml for 24 hours and biotin supplementation at 50 µM for 2 hours.

To test this hypothesis, we studied the ubiquitination of GLUL with or without pomalidomide. As mentioned previously, GLUL is an endogenous substrate of CRBN that, according to our results, is less ubiquitinated upon IMiD treatment. We performed a BioE3 experiment transiently transfecting BirA-CRBN and Clover-GLUL and treating the cells with pomalidomide and/or BTZ. We purified the biotinylated proteins by streptavidin beads and observed that GLUL is ubiquitinated by CRBN. Interestingly, the presence of pomalidomide decreased its ubiquitination, reinforcing our data (Fig. 5b; Supplementary file 1). This reduction in the ubiquitination was also confirmed by GFP-trap pull-down (Fig. 5c; Supplementary file 1). Accordingly, by confocal microscopy we see a decrease of biotinylated proteins co-localizing with Clover-GLUL in the cytoplasm, presumably corresponding to ubiquitinated Clover-GLUL (Fig. 5d).

## Discussion

BioE3 is a powerful technique to identify specific substrates of RING-and HECT-type Ub E3 ligases, while the complex CRLs, the biggest class of E3s and currently the main enzymes harnessed for the TPD strategies, have not yet been tested. Here we show that BioE3 can be applicable to CRLs, by identifying both endogenous targets and neosubstrates of the substrate receptor CRBN. We also proved its sensitivity to detect alterations of substrate specificity due to degrader treatment, i.e. IMiDs. These results are in agreement with those of Huang et al. (22), who applied a similar technique, E-STUB, to identify substrates of various molecular glues and PROTACs. Importantly, BioE3, as well as similar techniques recently developed (22, 23), can identify E3 substrates through ubiquitination status, independent of number of Ubs, Ub-chain type or reliance upon degradation by the UPS.

IMiDs are known to induce degradation of zinc-finger transcription factors in myeloma cells, like IKZF1 and IKZF3, and like SALL4 in embryonic stem cells, which is behind their therapeutic, but also teratogenic functions (40, 54). These drugs are currently approved for clinical use in different diseases, including multiple myeloma (9). However, the recruitment of new substrates by these drugs raises the question of the presence of unknown neosubstrates that could not only lead to undesired off-target effects, but that also could open new therapeutic opportunities in other diseases. We adapted BioE3 cells for an unbiased and systematic discovery of putative CRBN neosubstrates upon pomalidomide treatment. We did not identify the aforementioned neosubstrates, probably because HEK293FT express none or too little of those targets to be detected, underlining that the neosubstrate landscape may be different from one cell type to another.

Using BioE3 we were able to identify 133 putative neosubstrates and, among them, we found zinc-finger proteins, such as TRIM28 and ZBTB33. The neosubstrates were significantly enriched in RNA-binding proteins, which has also been recently observed by Baek *et al*. (51). Moreover, we validated CSDE1 as a neosubstrate of CRBN, an mRNA and stress-granule associated protein that has been linked to different types of cancer (52). In agreement with our data, CSDE1 was previously identified as a CRBN-AirID proximal interactor in HEK293FT cells upon pomalidomide treatment (36). In addition, we also found an enrichment in cytoskeletal proteins, like MYH9, which was recently described to be ubiquitinated by CRBN in response to IMiDs (37). This reinforces the conjecture of unknown therapeutic roles of IMiDs beyond multiple myeloma, but also raises questions about the unexpected secondary effects generated by the degradation of novel proteins. Notably, among the new putative neosubstrates presented here, some of them have features typical for thalidomide recognition, as the zinc finger or the G-loop, as is the case for KAISO (ZBTB33) or CSDE1, respectively (39, 53). However, we could not detect zinc fingers or G-loop motifs in all of them, suggesting that there might be other structural motifs key for the recognition by CRBN upon IMiDs treatment that remain hidden. In any case, BioE3 can be a powerful tool to define neosubstrates in the development of degraders, profiling secondary targets when a main target is desired, or identifying novel unexpected targets for a degrader that may have therapeutic use.

Even though a lot of effort has been dedicated to identifying neosubstrates of CRBN, not as much is known about the endogenous substrates of the substrate receptor. We tried to shed light to this question and identified the endogenous substrates of CRBN by two means. First, we assumed that ubiquitination by CRBN mainly promotes degradation of their target proteins by the UPS, and we identified substrate levels in absence or presence of the proteasomal inhibitor BTZ. Second, we assumed that CRBN works best when its cullin is NEDDylated, and we identified substrate levels in the absence or presence of the NEDDylation inhibitor MLN4924. We identified different subsets of putative substrates with the two treatments, with a significant overlap between them, but also with some proteins specific for each approach. This suggests that CRBN-mediated ubiquitination might not always lead to protein degradation, and that even in absence of NEDDylation, CRBN is proximal to some bioUb-modified proteins, perhaps preparing to extend a Ub chain. Although beyond the scope of this work, this extensive list of putative endogenous substrates could be of value in the discovery of shared degrons or motifs. Previously, Ichikawa et al. discovered C-terminal cyclic imides as a CRBN degron (55). Other features might still be unrevealed.

Among the putative substrates identified we found GLUL, one of the few endogenous substrates previously described (43). When considering all putative substrates, we found that CRBN is implicated in numerous cellular processes like cAMP and Wnt signaling pathways (46, 47), regulation of chloride channels (48), cell cycle, cell death and response to stress, among others. Thus, besides its role as a workhorse for TPD strategies, physiological roles for CRBN should be more carefully considered to assess secondary effects of TPD therapeutics.

## Conclusions

Molecular glues and PROTACs change the specificity of the substrate receptor, not only promoting neosubstrate degradation, but also affecting the ubiquitination of the endogenous substrates. In this work we observe a decrease in the ubiquitination of what we defined as the endogenous substrates of CRBN by BioE3 when treating the samples with pomalidomide, suggesting a competition for the binding with the neosubstrates. In fact, it was previously reported that IMiDs block the binding of some endogenous substrates to CRBN, like in the case of MEIS2 (56). Our results show that this is also true for GLUL. The rewiring of the different cellular functions of CRBN should be taken in consideration when developing TPD strategies, as it could lead to unexpected secondary effects, likely depending on the duration and concentration of drug treatment, as well as cell- or tissue-type. Our results highlight that E3-linked degraders like iMiDs can cause a significant rearrangement of the endogenous ubiquitination landscape, beyond the targeted reduction of the intended neosubstrate.

## List of abbreviations

bio^GEF^: Low affinity AviTag
BTZ: Bortezomib
CRBN: Cereblon
CRL: Cullin-RING E3 ligase
CRL4^CRBN^: CUL4–RBX1–DDB1–CRBN E3 complex
CSDE1: Cold Shock Domain Containing E1
DOX: Doxycycline
DUB: Deubiquitinating enzyme
GLUL: Glutamine Synthetase
HECT: Homology to E6AP C Terminus
IMiD: Immunomodulatory Imide Drug
LC-MS/MS: Liquid chromatography-mass spectrometry
MG: Molecular Glue
PMF: Potential Mean Force
PROTAC: Proteolysis Targeting Chimeras
RBR: RING-Between-RING
RING: Really Interesting New Gene
SALL4: Spalt-like 4
TPD: Targeted Protein Degradation
Ub: Ubiquitin
UbL: Ubiquitin-like
UPS: Ubiquitin Proteasome System

## Declarations

### Ethics approval and consent to participate

Not applicable

### Consent for publication

Not applicable

### Availability of data and materials

The datasets generated and/or analyzed during the current study are available in the ProteomeXchange Consortium via the PRIDE partner repository(57) with the dataset identifier PXD055877. All the other data generated or analyzed during this study are included in this published article [and its supplementary information files].

### Competing interests

C.G. is co-founder and Drug Discovery Scientific Advisor of Oniria Therapeutics. The other authors declare that they have no competing interests.

## Funding

R.B. and J.D.S. were funded by MCIN/AEI/10.13039/501100011033, projects PID2023-147399NB-I00 and PID2020-114178GB-I00 and CEX2021-001136-S and CEX2021-001202-M Severo Ochoa Excellence Program and additional support provided by the Diputación Foral de Bizkaia, Programa Transferencia Tecnológica 2023, project 6/12/TT/2023 /00001, and the Department of Industry, Tourism, and Trade of the Basque Country Government (Elkartek Research Programs); J.J.-J. was funded by grant CNS2022-135307 funded by the MCIN/AEI/10.13039/501100011033 and the European Union NextGeneration EU/PRTR; V.M., C.G.-P. and M.P.-R. were funded by FPI grants funded by MCIN/AEI /10.13039/501100011033 and FSE, PRE2018-086230, PRE2021-099359 and PRE2022-104553, respectively; L.M.-C. was funded by FPU grant FPU20/05282 (funded by Ministerio de Universidades para la Formación de Profesorado Universitario). O.B.-G., J.D.S. and R.B. acknowledge funding by the grant 765445-EU (UbiCODE Program). U.M. was funded by the Spanish MCIU (PID2020-117333GB-I00 (FEDER/EU)). Access to HPC facilities were granted through the Red Española de Supercomputación (BCV-2021-3-0006 and BCV-2021-2-0005).

## Authors’ contributions

L.M.-C., O.B.-G., R.B. and J.D.S. conceived and designed the work; L.M.-C., O.B.-G., M.P.-R., V.M., C.G.-P., A.S., C.P., A.U., M.A., I.I. and F.E. acquired and analyzed the data; L.M.-C., O.B.-G., C.G., J.J.-J., U.M., R.B. and J.D.S. interpreted the data; L.M.-C., M.P.-R. and R.B. drafted the manuscript; and J.D.S., O.B.-G. and C.G., J.J.-J. substantively revised it. All authors read and approved the final manuscript.

## Acknowledgements

R.B., J.D.S., U.M. and C.G. acknowledge networking support from the European Cooperation for Science & Technology ProteoCure COST Action (CA20113). C.G. and J.J.-J. are members of the Computational Biology Drug Design Consolidated Research Group supported by the Generalitat de Catalunya (2021SGR00671).

## Supplementary files

**Fig. S1. Images of uncropped Western blot membranes.** The parts of the blots used in the Figures are delimited by dotted blue lines. Membranes with molecular weight markers are shown to the right of each blot.

**Fig. S2. Optimization of the experimental conditions for CRBN BioE3. a, b, c)** Western blot of BioE3 experiments performed on HEK293FT stable cell lines expressing TRIPZ-bio^GEF^Ub or TRIPZ-bio^GEF^Ubnc and transfected with EFS-BirA-CRBN, EFS-CRBN-BirA or EFS-BirA-CRBN^W386A^. Indicated samples were induced with doxycycline (DOX) at 1 µg/ml for 24 hours, treated with 200 nM bortezomib (BTZ) for 6 hours, 1 µM MLN4924 for 24 hours or 10 µM pomalidomide (Poma) for 6 hours and supplemented with 50 µM biotin for 2 hours. Molecular weight markers are shown to the left of the blots in kDa, antibodies used are indicated to the right. All BioE3 experiments were performed by pre-incubating the cells in dialyzed FBS-containing media prior to transfections.

**Fig. S3. STRING network analysis of CRBN ubiquitinated substrates.** Substrates defined in Figure 3b show a highly interconnected network composed of 71% of the proteins. Highly interconnected sub-clusters were derived and characterized using MCODE. Color, transparency and size of the nodes were discretely mapped to the Log2 enrichment value as indicated.

**Fig. S4. Gene ontology scatterplot of the CRBN ubiquitinated substrates.** REVIGO plots were generated for targets defined in Figure 3b. Colors indicate the -Log10 p-value as shown in the Figure, and size of the bubble indicates the size of each term. Only terms with a p-value < 0.05 are represented.

**Fig. S5. STRING network analysis of CRBN NEDDylation-dependent substrates.** Substrates defined in Figure 3c show a highly interconnected network composed of 64% of the proteins. Highly interconnected sub-clusters were derived and characterized using MCODE. Color, transparency and size of the nodes were discretely mapped to the Log2 enrichment value as indicated.

**Fig. S6. Gene ontology scatterplot of CRBN NEDDylation-dependent substrates.** REVIGO plots were generated for targets defined in Figure 3c. Colors indicate the -Log10 p-value as shown in the Figure, and size of the bubble indicates the size of each term. Only terms with a p-value < 0.05 are represented.

**Fig. S7. STRING network analysis of CRBN neosubstrates.** The neosubstrates upon pomalidomide treatment defined in Figure 4a show a highly interconnected network composed of 71% of the proteins. Highly interconnected sub-clusters were derived and characterized using MCODE. Color, transparency and size of the nodes were discretely mapped to the Log2 enrichment value as indicated.

**Fig. S8. Gene ontology scatterplot of the neosubstrates of CRBN.** REVIGO plots were generated for targets defined in Figure 4a. Colors indicate the -Log10 p-value as shown in the Figure, and size of the bubble indicates the size of each term. Only terms with a p-value < 0.05 are represented.

**Supplementary Table 1.** LC-MS/MS processed data of CRBN BioE3 treated or not with proteasomal inhibitor bortezomib (sheet 1). Gene Ontology analysis of CRBN targets (sheet 2) and selected GO terms by Revigo (sheet 3).

**Supplementary Table 2.** LC-MS/MS processed data of CRBN BioE3 with proteasomal inhibitor bortezomib, with or without NEDDylation inhibitor MLN4924 (sheet 1). Gene Ontology analysis of CRBN targets (sheet 2) and selected GO terms by Revigo (sheet 3).

**Supplementary Table 3.** LC-MS/MS processed data of CRBN BioE3 with proteasomal inhibitor bortezomib, with or without pomalidomide treatment (sheet 1). Gene Ontology analysis of CRBN targets (sheet 2) and selected GO terms by Revigo (sheet 3).

## References

1. Dikic I, Schulman BA. An expanded lexicon for the ubiquitin code. Nature Reviews Molecular Cell Biology. 2023;24(4):273–87.

2. Cappadocia L, Lima CD. Ubiquitin-like Protein Conjugation: Structures, Chemistry, and Mechanism. Chem Rev. 2018;118(3):889–918.

3. Lange SM, Armstrong LA, Kulathu Y. Deubiquitinases: From mechanisms to their inhibition by small molecules. Mol Cell. 2022;82(1):15–29.

4. Zheng N, Shabek N. Ubiquitin Ligases: Structure, Function, and Regulation. Annu Rev Biochem. 2017;86:129–57.

5. Morreale FE, Walden H. Types of Ubiquitin Ligases. Cell. 2016;165(1):248–e1.

6. Baek K, Scott DC, Schulman BA. NEDD8 and ubiquitin ligation by cullin-RING E3 ligases. Curr Opin Struct Biol. 2021;67:101–9.

7. Ito T, Ando H, Suzuki T, Ogura T, Hotta K, Imamura Y, et al. Identification of a primary target of thalidomide teratogenicity. Science. 2010;327(5971):1345–50.

8. Oleinikovas V, Gainza P, Ryckmans T, Fasching B, Thoma NH. From Thalidomide to Rational Molecular Glue Design for Targeted Protein Degradation. Annu Rev Pharmacol Toxicol. 2024;64:291–312.

9. Tsai JM, Nowak RP, Ebert BL, Fischer ES. Targeted protein degradation: from mechanisms to clinic. Nat Rev Mol Cell Biol. 2024;25(9):740–57.

10. Belcher BP, Ward CC, Nomura DK. Ligandability of E3 Ligases for Targeted Protein Degradation Applications. Biochemistry. 2023;62(3):588–600.

11. Roux KJ, Kim DI, Raida M, Burke B. A promiscuous biotin ligase fusion protein identifies proximal and interacting proteins in mammalian cells. J Cell Biol. 2012;196(6):801–10.

12. Coyaud E, Mis M, Laurent EM, Dunham WH, Couzens AL, Robitaille M, et al. BioID-based Identification of Skp Cullin F-box (SCF)beta-TrCP1/2 E3 Ligase Substrates. Mol Cell Proteomics. 2015;14(7):1781–95.

13. Zhang H, Li S, Liu P, Lee FHF, Wong AHC, Liu F. Proteomic analysis of the cullin 4B interactome using proximity-dependent biotinylation in living cells. Proteomics. 2017;17(8).

14. O’Connor HF, Lyon N, Leung JW, Agarwal P, Swaim CD, Miller KM, et al. Ubiquitin-Activated Interaction Traps (UBAITs) identify E3 ligase binding partners. EMBO Rep. 2015;16(12):1699–712.

15. Kumar R, Gonzalez-Prieto R, Xiao Z, Verlaan-de Vries M, Vertegaal ACO. The STUbL RNF4 regulates protein group SUMOylation by targeting the SUMO conjugation machinery. Nat Commun. 2017;8(1):1809.

16. Salas-Lloret D, Jansen NS, Nagamalleswari E, van der Meulen C, Gracheva E, de Ru AH, et al. SUMO-activated target traps (SATTs) enable the identification of a comprehensive E3-specific SUMO proteome. Sci Adv. 2023;9(31):eadh2073.

17. Watanabe M, Saeki Y, Takahashi H, Ohtake F, Yoshida Y, Kasuga Y, et al. A substrate-trapping strategy to find E3 ubiquitin ligase substrates identifies Parkin and TRIM28 targets. Commun Biol. 2020;3(1):592.

18. Ramirez J, Lectez B, Osinalde N, Sivá M, Elu N, Aloria K, et al. Quantitative proteomics reveals neuronal ubiquitination of Rngo/Ddi1 and several proteasomal subunits by Ube3a, accounting for the complexity of Angelman syndrome. Hum Mol Genet. 2018;27(11):1955–71.

19. Alduntzin U, Lectez B, Presa N, Osinalde N, Fernandez M, Elu N, et al. Angelman Syndrome causing UBE3A ligase displays predominantly synaptic ubiquitination activity in the mouse brain. Research Square. 2023.

20. Barroso-Gomila O, Merino-Cacho L, Muratore V, Perez C, Taibi V, Maspero E, et al. BioE3 identifies specific substrates of ubiquitin E3 ligases. Nat Commun. 2023;14(1):7656.

21. Fernandez-Suarez M, Chen TS, Ting AY. Protein-protein interaction detection in vitro and in cells by proximity biotinylation. J Am Chem Soc. 2008;130(29):9251–3.

22. Huang HT, Lumpkin RJ, Tsai RW, Su S, Zhao X, Xiong Y, et al. Ubiquitin-specific proximity labeling for the identification of E3 ligase substrates. Nat Chem Biol. 2024.

23. Mukhopadhyay U, Levantovsky S, Carusone TM, Gharbi S, Stein F, Behrends C, et al. A ubiquitin-specific, proximity-based labeling approach for the identification of ubiquitin ligase substrates. Sci Adv. 2024;10(32):eadp3000.

24. Tyanova S, Temu T, Sinitcyn P, Carlson A, Hein MY, Geiger T, et al. The Perseus computational platform for comprehensive analysis of (prote)omics data. Nat Methods. 2016;13(9):731–40.

25. Snel B, Lehmann G, Bork P, Huynen MA. STRING: a web-server to retrieve and display the repeatedly occurring neighbourhood of a gene. Nucleic Acids Res. 2000;28(18):3442–4.

26. Shannon P, Markiel A, Ozier O, Baliga NS, Wang JT, Ramage D, et al. Cytoscape: a software environment for integrated models of biomolecular interaction networks. Genome Res. 2003;13(11):2498–504.

27. Bader GD, Hogue CW. An automated method for finding molecular complexes in large protein interaction networks. BMC Bioinformatics. 2003;4:2.

28. Reimand J, Arak T, Adler P, Kolberg L, Reisberg S, Peterson H, et al. g:Profiler-a web server for functional interpretation of gene lists (2016 update). Nucleic Acids Res. 2016;44(W1):W83–9.

29. Supek F, Bosnjak M, Skunca N, Smuc T. REVIGO summarizes and visualizes long lists of gene ontology terms. PLoS One. 2011;6(7):e21800.

30. Heberle H, Meirelles GV, da Silva FR, Telles GP, Minghim R. InteractiVenn: a web-based tool for the analysis of sets through Venn diagrams. BMC Bioinformatics. 2015;16(1):169.

31. Frisch MJ, Trucks GW, Schlegel HB, Scuseria GE, Robb MA, Cheeseman JR, et al. Gaussian 16 Rev. C.01. Wallingford, CT2016.

32. Peters MB, Yang Y, Wang B, Fusti-Molnar L, Weaver MN, Merz KM, Jr. Structural Survey of Zinc Containing Proteins and the Development of the Zinc AMBER Force Field (ZAFF). J Chem Theory Comput. 2010;6(9):2935–47.

33. Miñarro-Lleonar M, Bertran-Mostazo A, Duro J, Barril X, Juarez-Jimenez J. Lenalidomide Stabilizes Protein-Protein Complexes by Turning Labile Intermolecular H-Bonds into Robust Interactions. J Med Chem. 2023;66(9):6037–46.

34. Case DA, Aktulga HM, Belfon K, Cerutti DS, Cisneros GA, Cruzeiro VWD, et al. AmberTools. J Chem Inf Model. 2023;63(20):6183–91.

35. Jarzynski C. Nonequilibrium Equality for Free Energy Differences. Physical Review Letters. 1997;78(14):2690–3.

36. Yamanaka S, Horiuchi Y, Matsuoka S, Kido K, Nishino K, Maeno M, et al. A proximity biotinylation-based approach to identify protein-E3 ligase interactions induced by PROTACs and molecular glues. Nat Commun. 2022;13(1):183.

37. Costacurta M, Sandow JJ, Maher B, Susanto O, Vervoort SJ, Devlin JR, et al. Mapping the IMiD-dependent cereblon interactome using BioID-proximity labelling. FEBS J. 2024.

38. Matyskiela ME, Couto S, Zheng X, Lu G, Hui J, Stamp K, et al. SALL4 mediates teratogenicity as a thalidomide-dependent cereblon substrate. Nat Chem Biol. 2018;14(10):981–7.

39. Sievers QL, Petzold G, Bunker RD, Renneville A, Slabicki M, Liddicoat BJ, et al. Defining the human C2H2 zinc finger degrome targeted by thalidomide analogs through CRBN. Science. 2018;362(6414).

40. Donovan KA, An J, Nowak RP, Yuan JC, Fink EC, Berry BC, et al. Thalidomide promotes degradation of SALL4, a transcription factor implicated in Duane Radial Ray syndrome. Elife. 2018;7.

41. Matyskiela ME, Lu G, Ito T, Pagarigan B, Lu CC, Miller K, et al. A novel cereblon modulator recruits GSPT1 to the CRL4(CRBN) ubiquitin ligase. Nature. 2016;535(7611):252–7.

42. Thul PJ, Akesson L, Wiking M, Mahdessian D, Geladaki A, Ait Blal H, et al. A subcellular map of the human proteome. Science. 2017;356(6340).

43. Nguyen TV, Lee JE, Sweredoski MJ, Yang SJ, Jeon SJ, Harrison JS, et al. Glutamine Triggers Acetylation-Dependent Degradation of Glutamine Synthetase via the Thalidomide Receptor Cereblon. Mol Cell. 2016;61(6):809–20.

44. Hjerpe R, Thomas Y, Chen J, Zemla A, Curran S, Shpiro N, et al. Changes in the ratio of free NEDD8 to ubiquitin triggers NEDDylation by ubiquitin enzymes. Biochem J. 2012;441(3):927–36.

45. Hjerpe R, Thomas Y, Kurz T. NEDD8 overexpression results in neddylation of ubiquitin substrates by the ubiquitin pathway. J Mol Biol. 2012;421(1):27–9.

46. Lee KM, Yang SJ, Choi JH, Park CS. Functional effects of a pathogenic mutation in Cereblon (CRBN) on the regulation of protein synthesis via the AMPK-mTOR cascade. J Biol Chem. 2014;289(34):23343–52.

47. Shen C, Nayak A, Neitzel LR, Adams AA, Silver-Isenstadt M, Sawyer LM, et al. The E3 ubiquitin ligase component, Cereblon, is an evolutionarily conserved regulator of Wnt signaling. Nat Commun. 2021;12(1):5263.

48. Chen YA, Peng YJ, Hu MC, Huang JJ, Chien YC, Wu JT, et al. The Cullin 4A/B-DDB1-Cereblon E3 Ubiquitin Ligase Complex Mediates the Degradation of CLC-1 Chloride Channels. Sci Rep. 2015;5:10667.

49. Eichner R, Heider M, Fernandez-Saiz V, van Bebber F, Garz AK, Lemeer S, et al. Immunomodulatory drugs disrupt the cereblon-CD147-MCT1 axis to exert antitumor activity and teratogenicity. Nat Med. 2016;22(7):735–43.

50. Schulze-Niemand E, Naumann M. The COP9 signalosome: A versatile regulatory hub of Cullin-RING ligases. Trends Biochem Sci. 2023;48(1):82–95.

51. Baek K, Metivier RJ, Roy Burman SS, Bushman JW, Lumpkin RJ, Abeja DM, et al. Unveiling the hidden interactome of CRBN molecular glues with chemoproteomics. bioRxiv. 2024:2024.09.11.612438.

52. Ciocia A, Mestre-Farras N, Vicent-Nacht I, Guitart T, Gebauer F. CSDE1: a versatile regulator of gene expression in cancer. NAR Cancer. 2024;6(2):zcae014.

53. Petzold G, Fischer ES, Thoma NH. Structural basis of lenalidomide-induced CK1alpha degradation by the CRL4(CRBN) ubiquitin ligase. Nature. 2016;532(7597):127–30.

54. Kronke J, Udeshi ND, Narla A, Grauman P, Hurst SN, McConkey M, et al. Lenalidomide causes selective degradation of IKZF1 and IKZF3 in multiple myeloma cells. Science. 2014;343(6168):301–5.

55. Ichikawa S, Flaxman HA, Xu W, Vallavoju N, Lloyd HC, Wang B, et al. The E3 ligase adapter cereblon targets the C-terminal cyclic imide degron. Nature. 2022;610(7933):775–82.

56. Fischer ES, Bohm K, Lydeard JR, Yang H, Stadler MB, Cavadini S, et al. Structure of the DDB1-CRBN E3 ubiquitin ligase in complex with thalidomide. Nature. 2014;512(7512):49–53.

57. Vizcaino JA, Csordas A, del-Toro N, Dianes JA, Griss J, Lavidas I, et al. 2016 update of the PRIDE database and its related tools. Nucleic Acids Res. 2016;44(D1):D447–56.

