## Supplementary figures and images for "Cullin-RING ligase BioE3 reveals molecular-glue-induced neosubstrates and rewiring of the endogenous Cereblon ubiquitome"

### Supplementary figure_Fig_S1

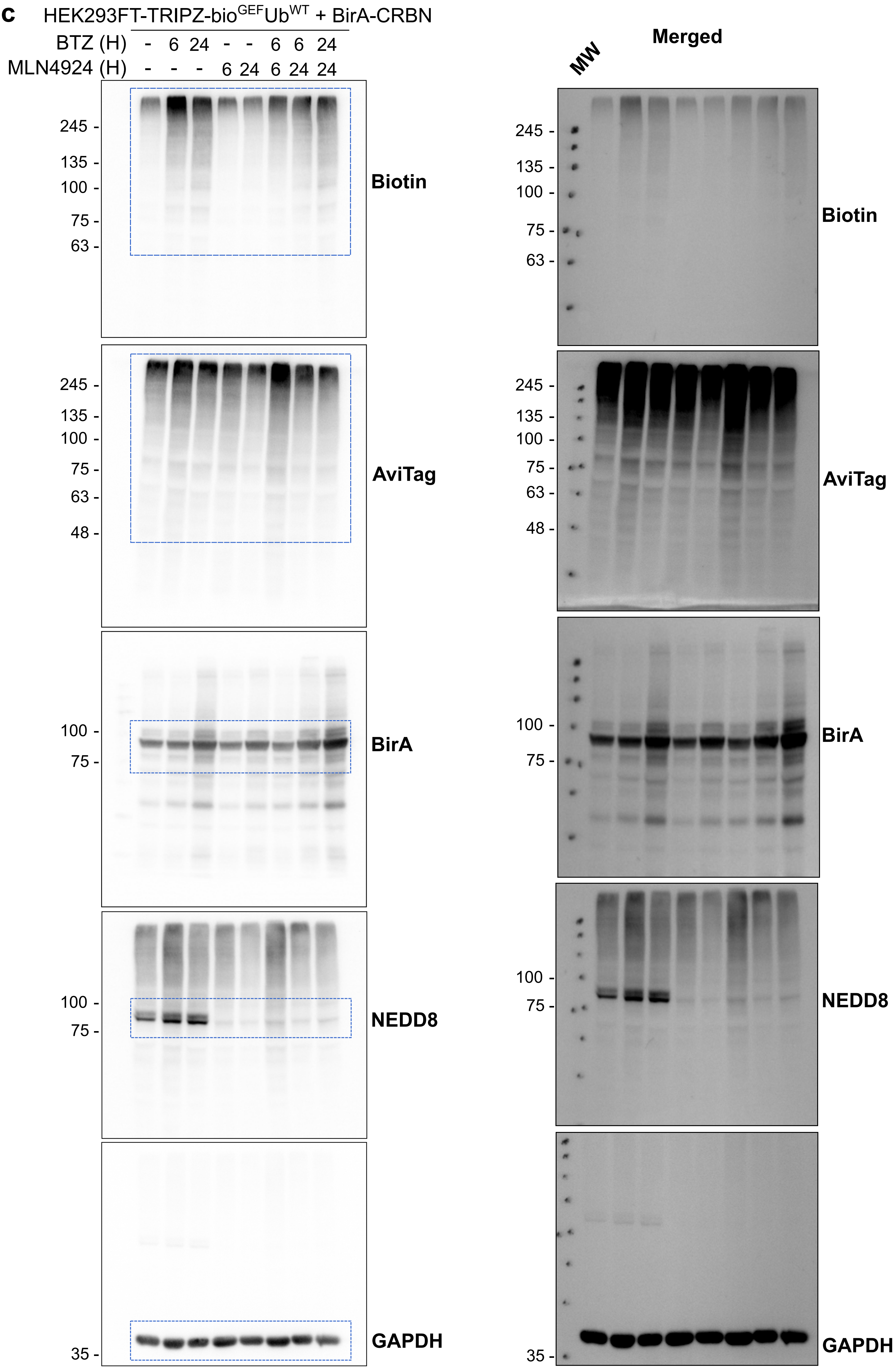

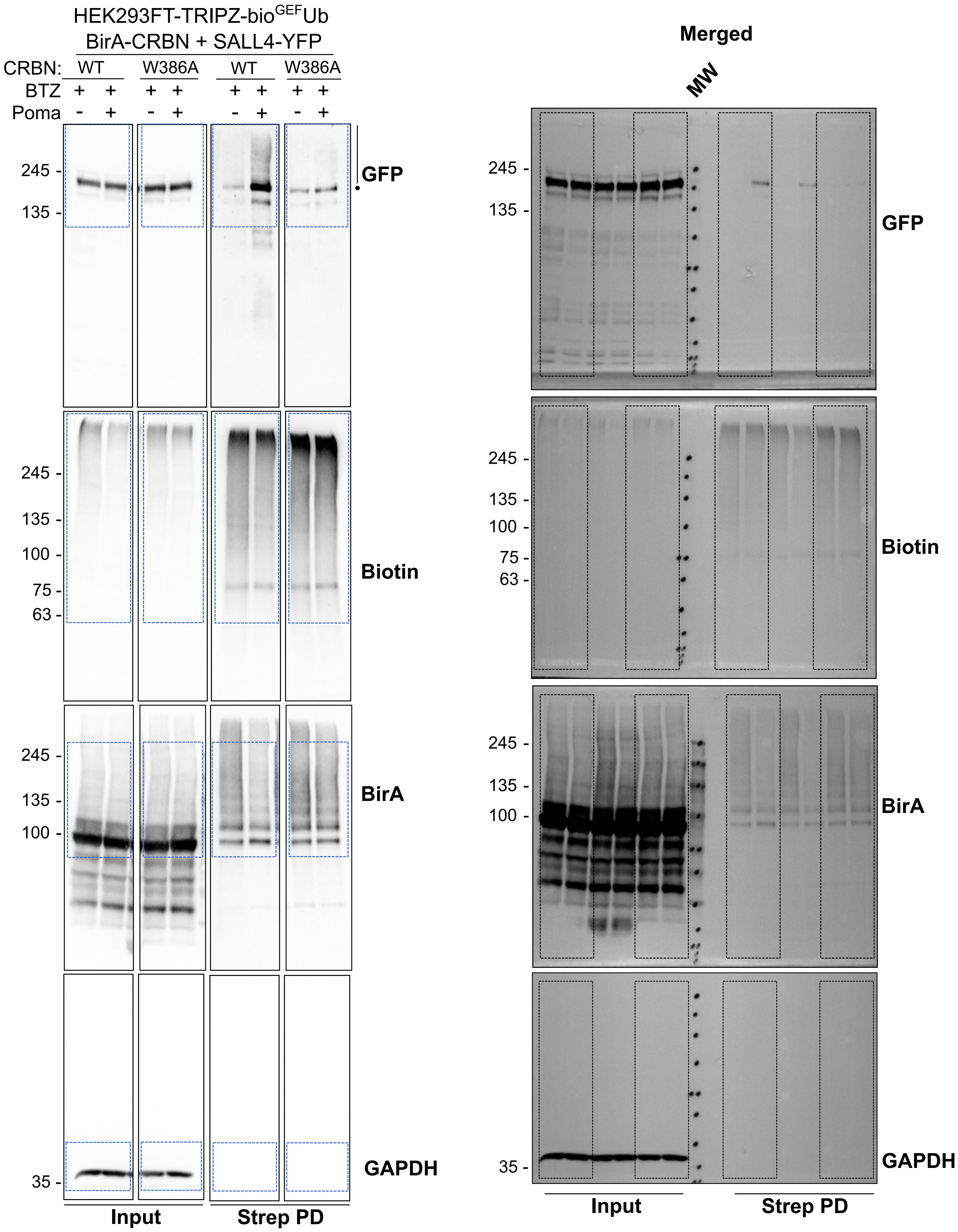

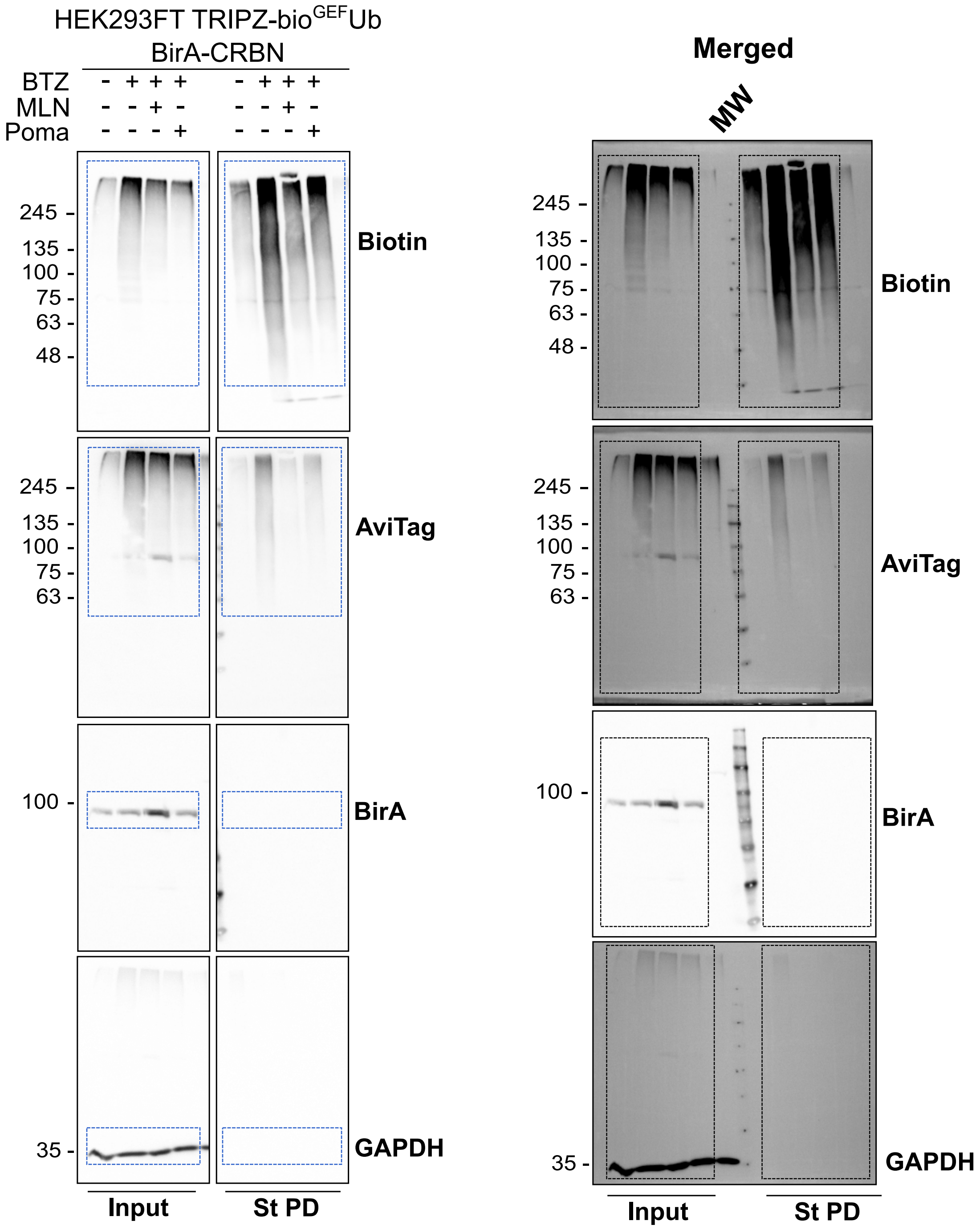

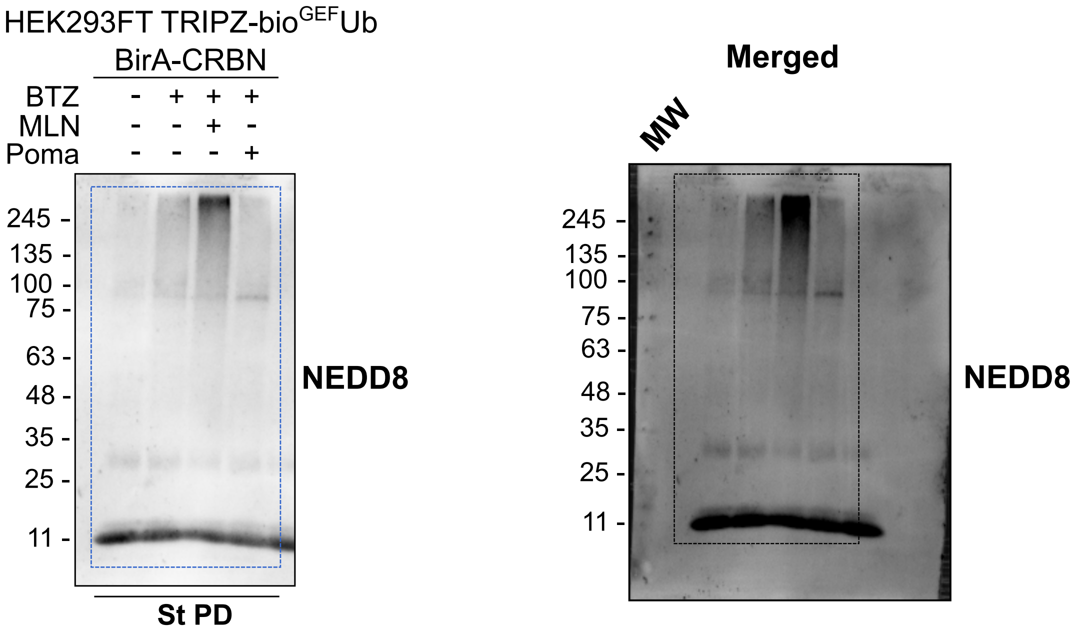

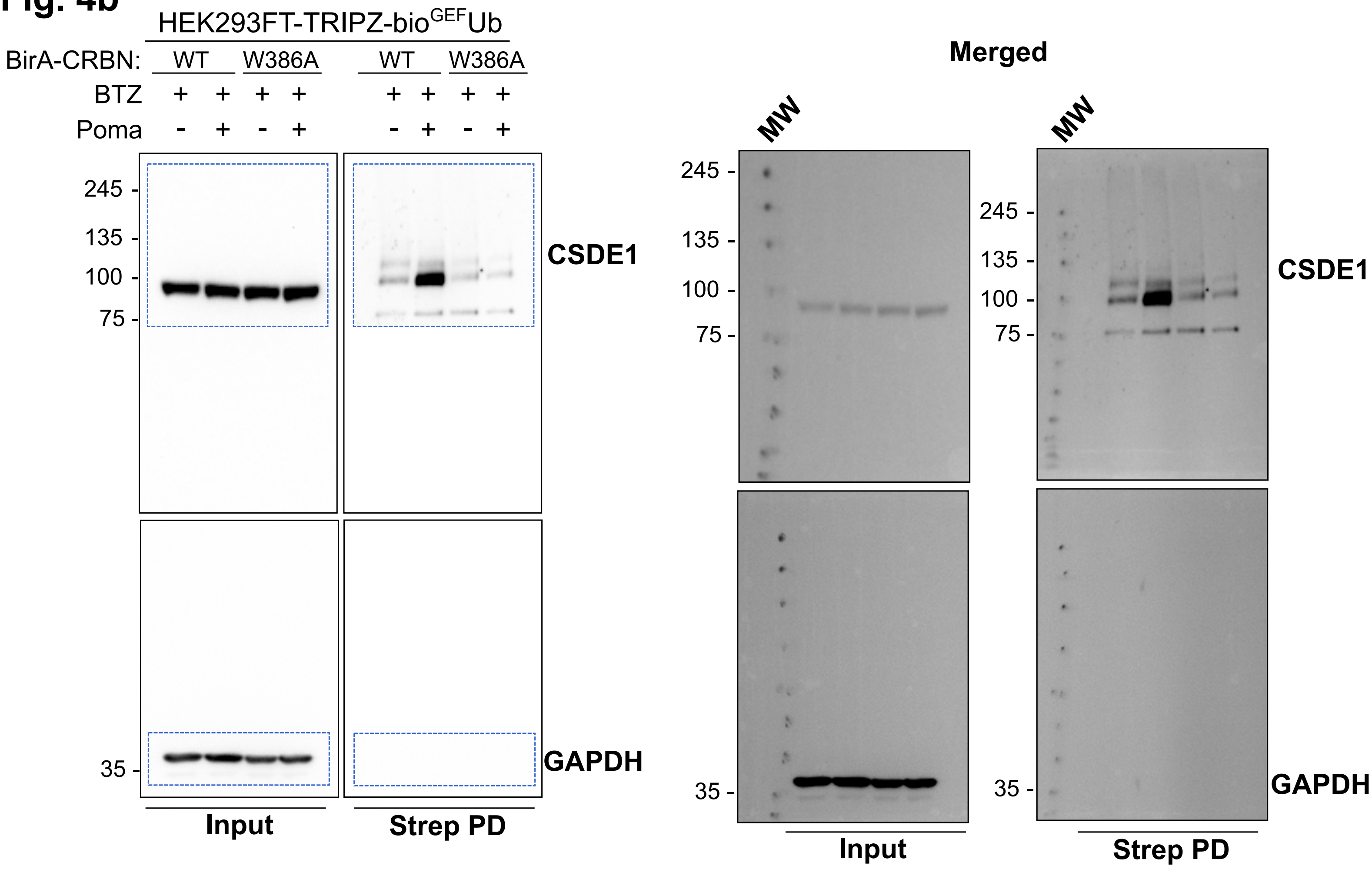

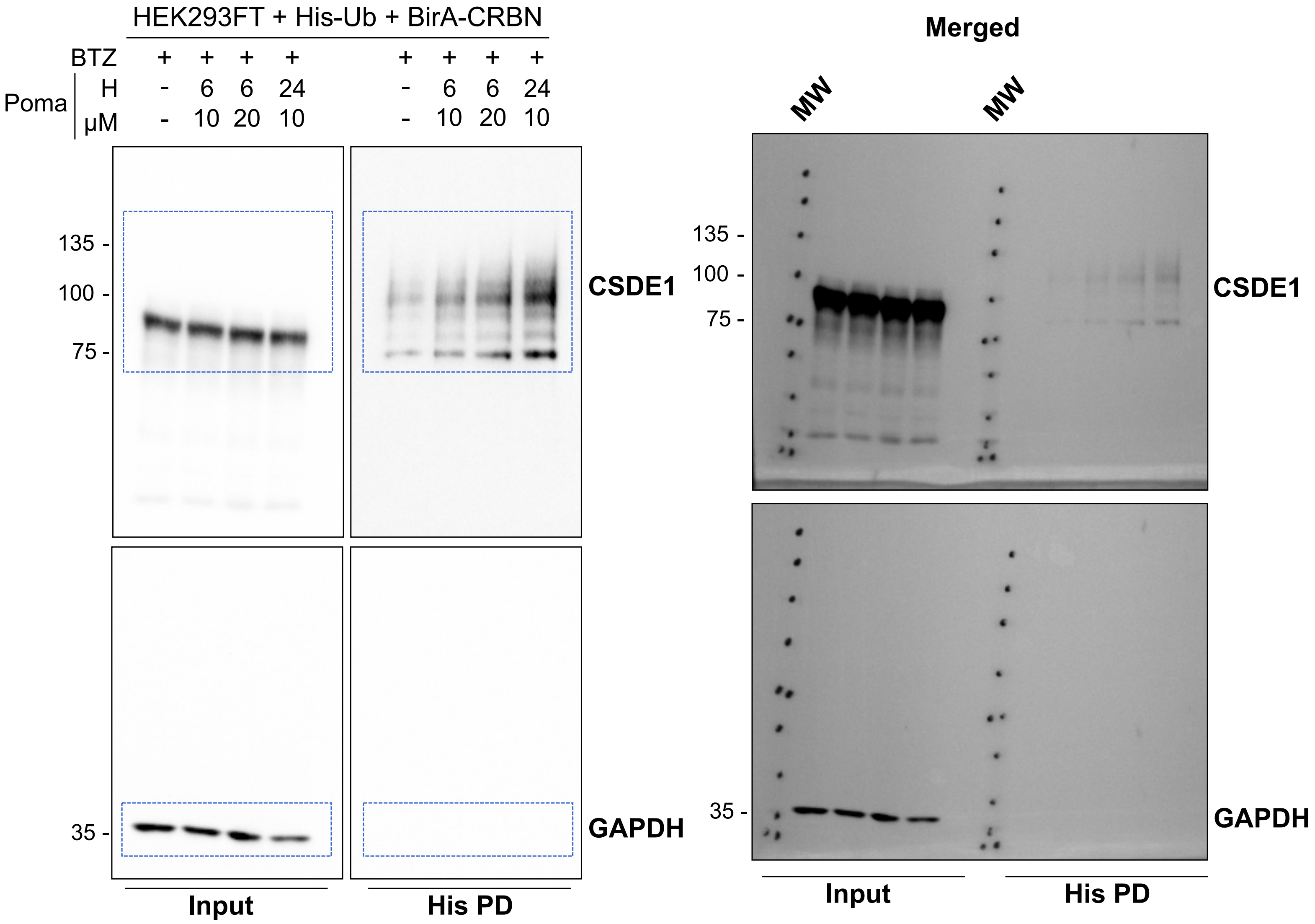

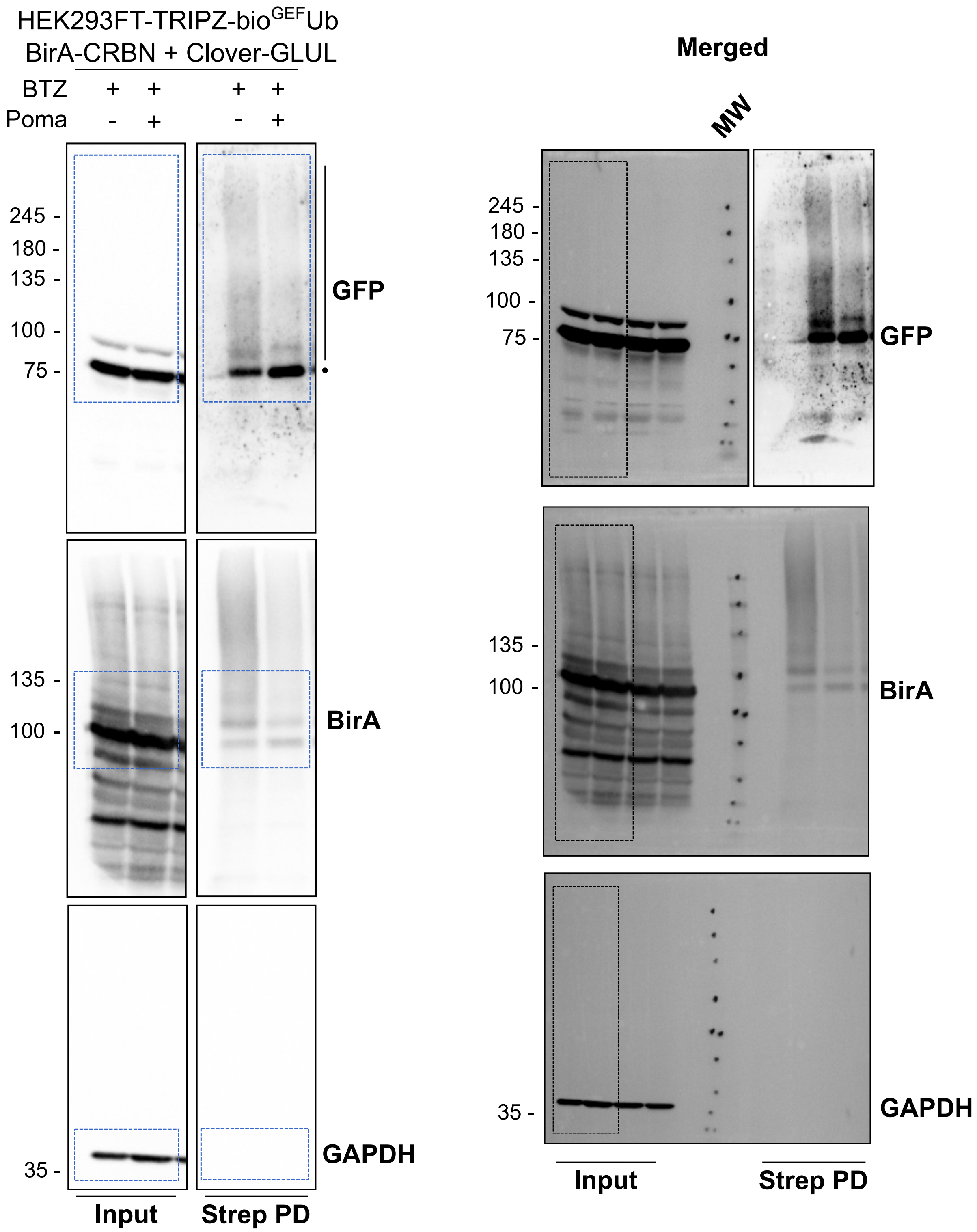

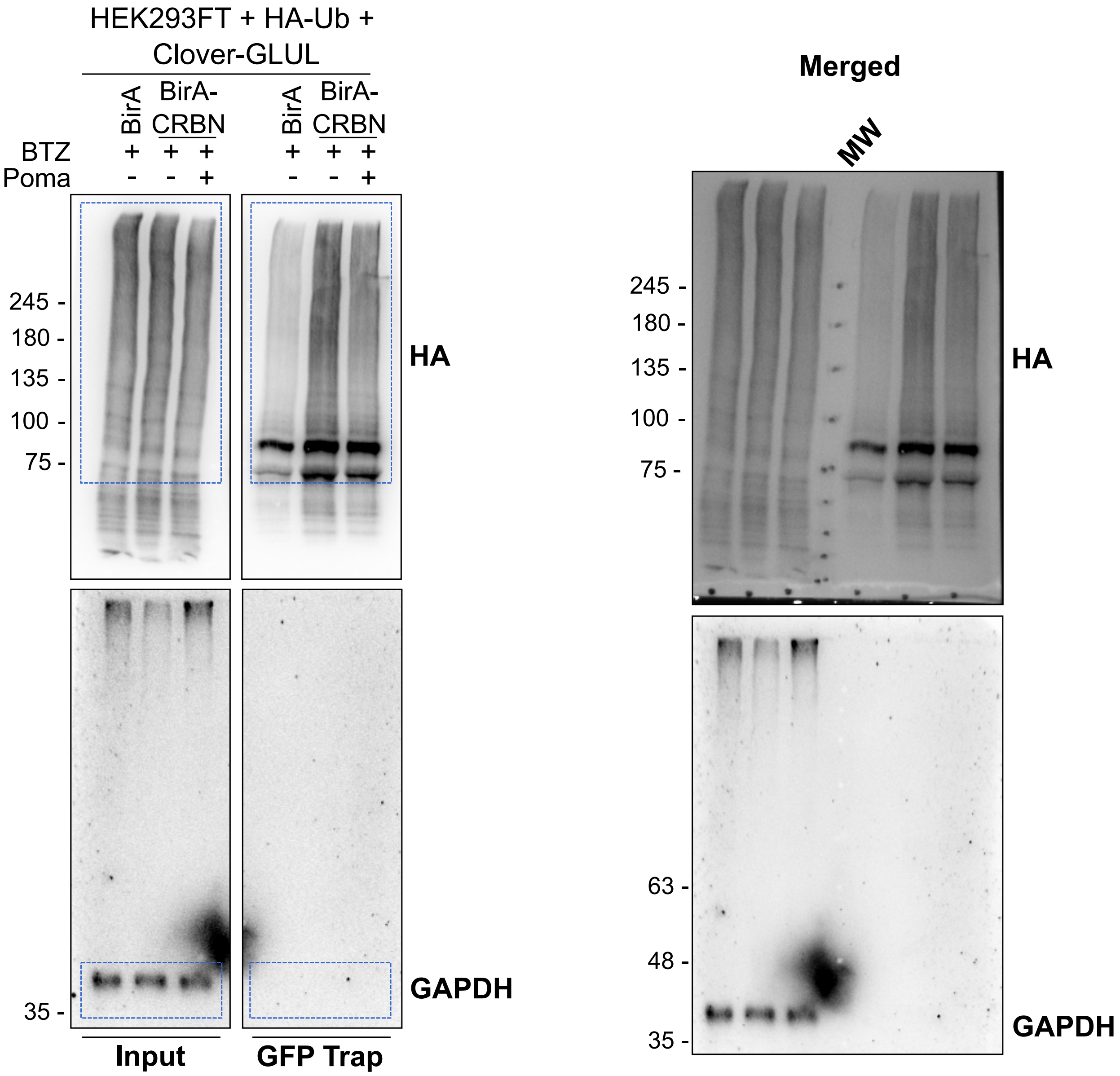

##### Supplementary Fig. 1a

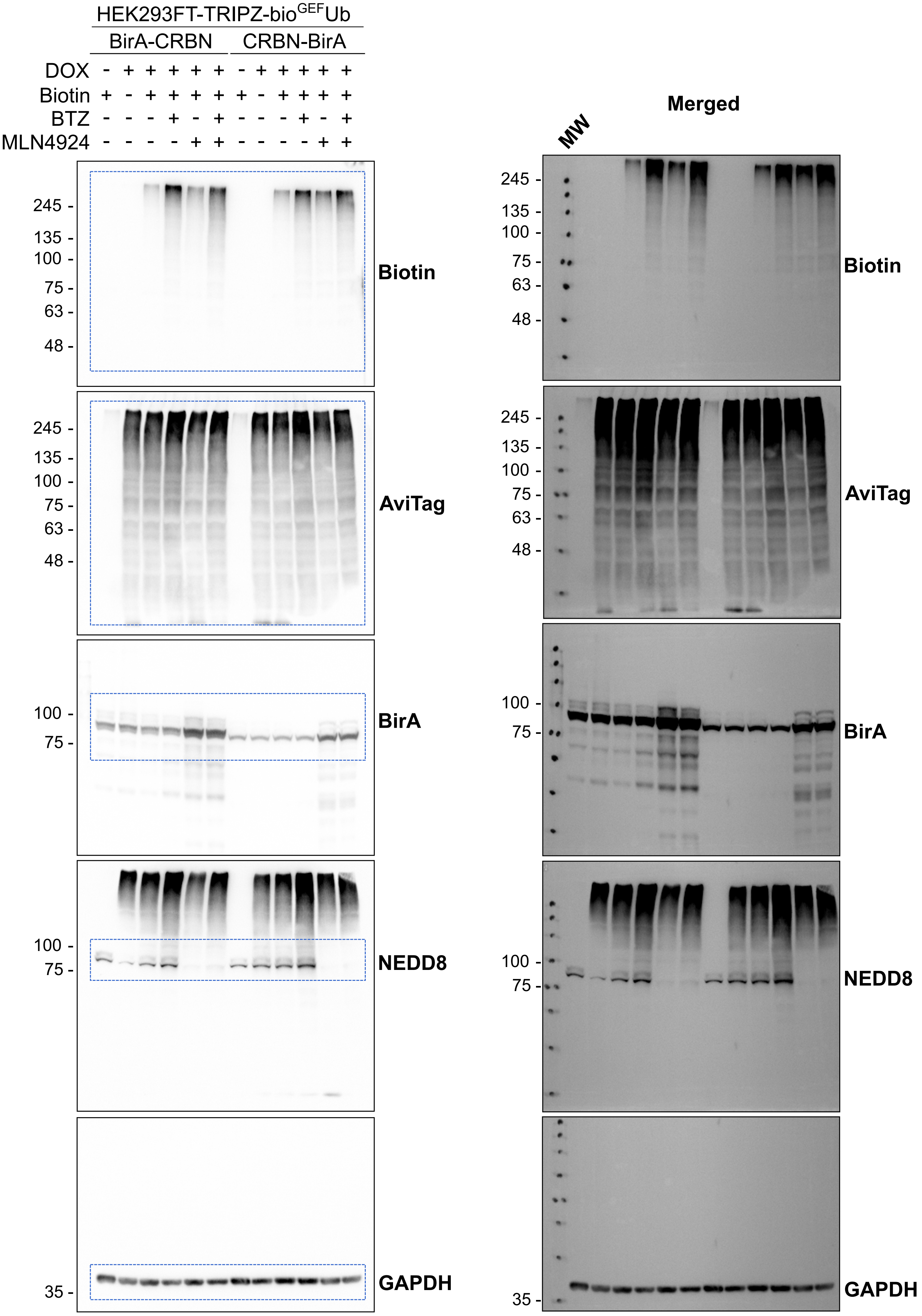

#### Supplementary Fig. 1b

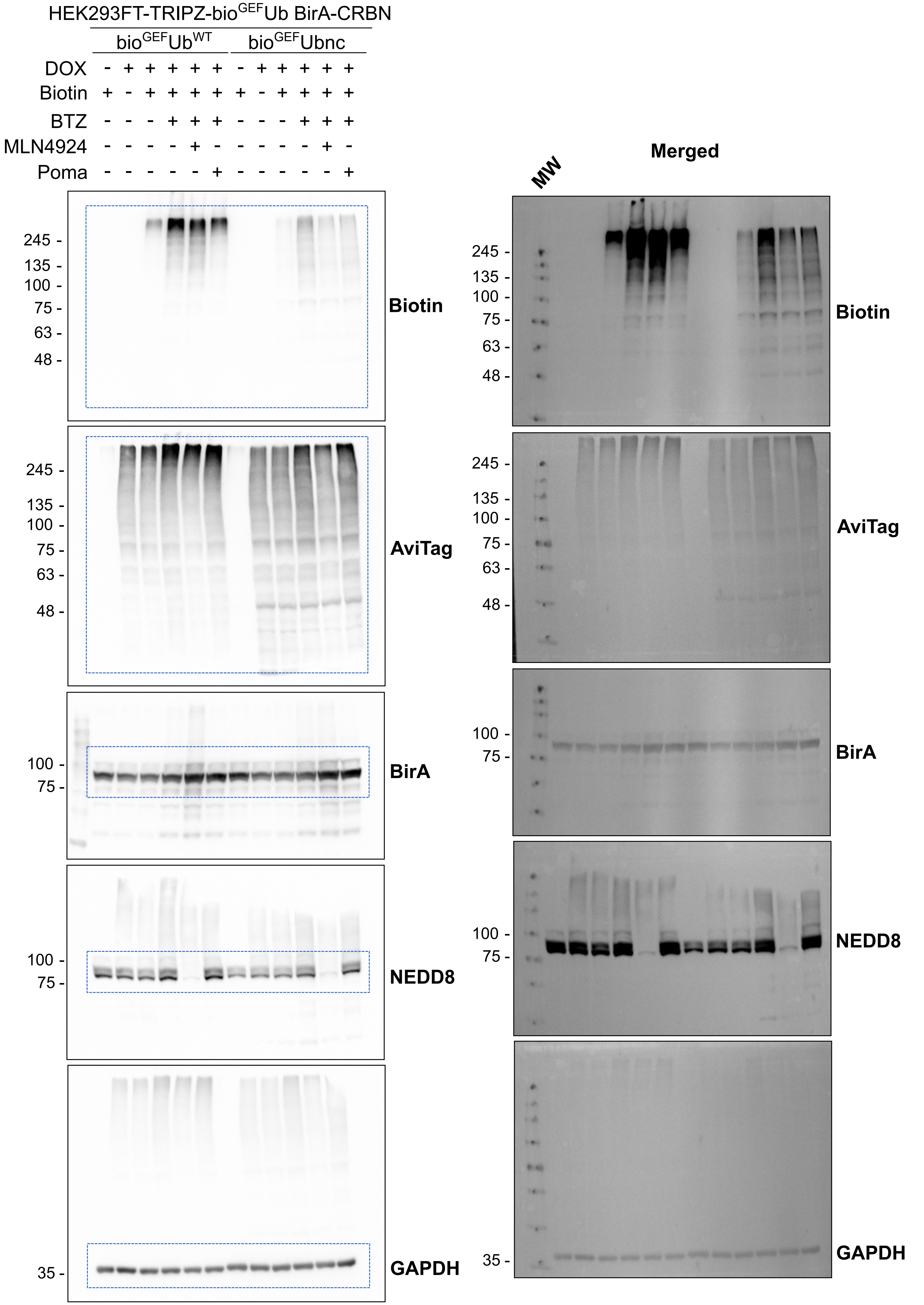

#### Supplementary Fig. 1c

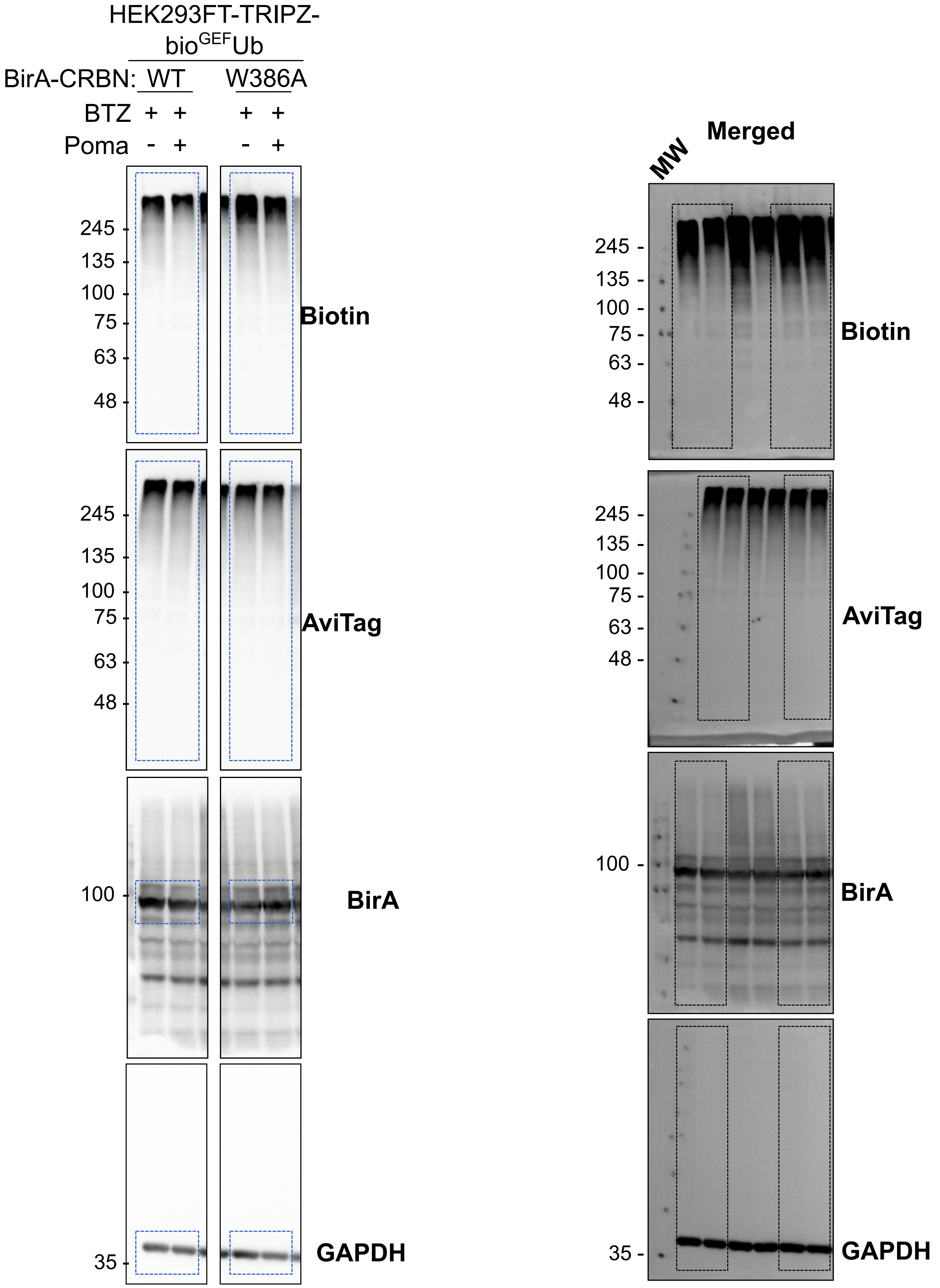

### Supplementary figure_Fig_S2

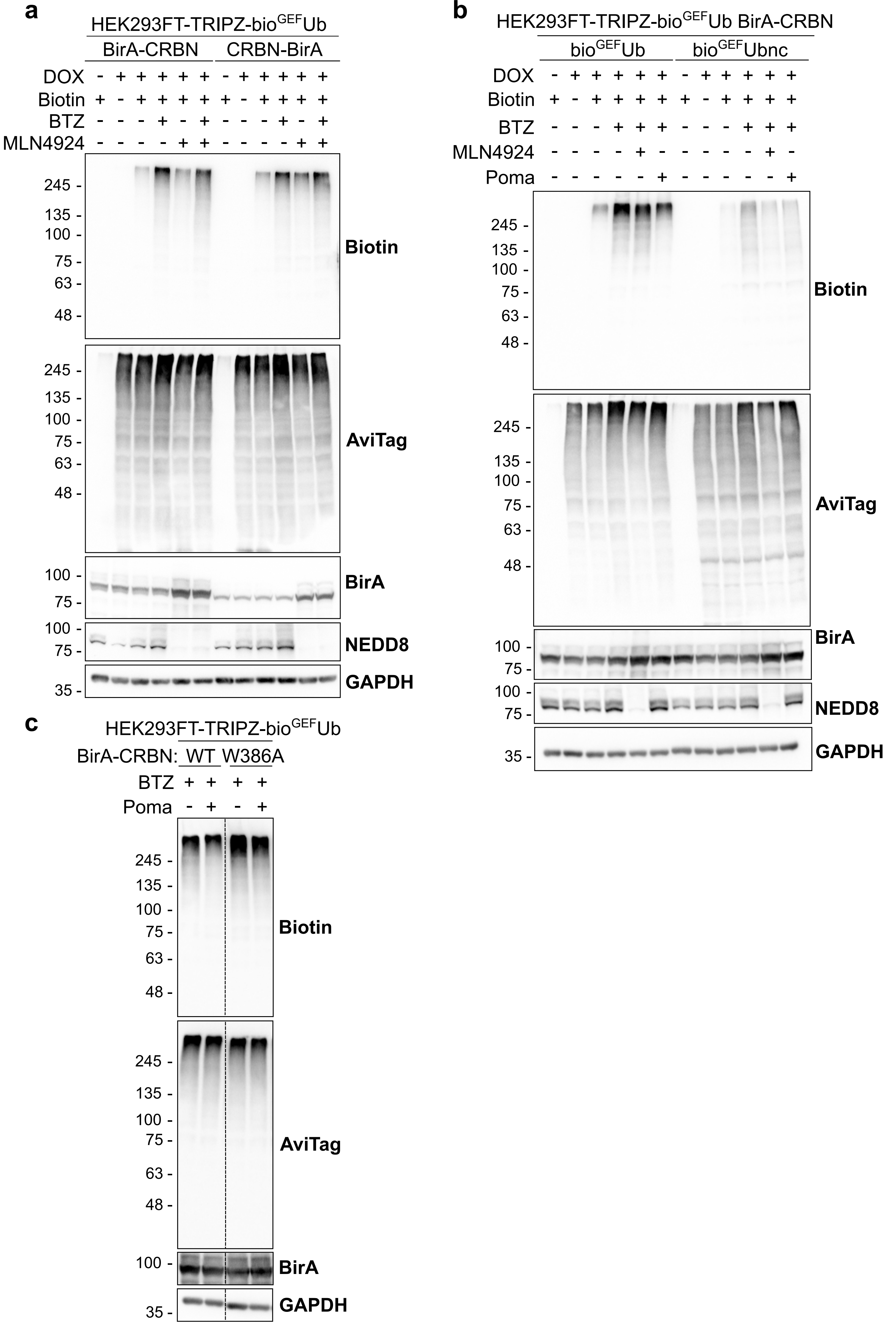

### Supplementary figure_Fig_S3

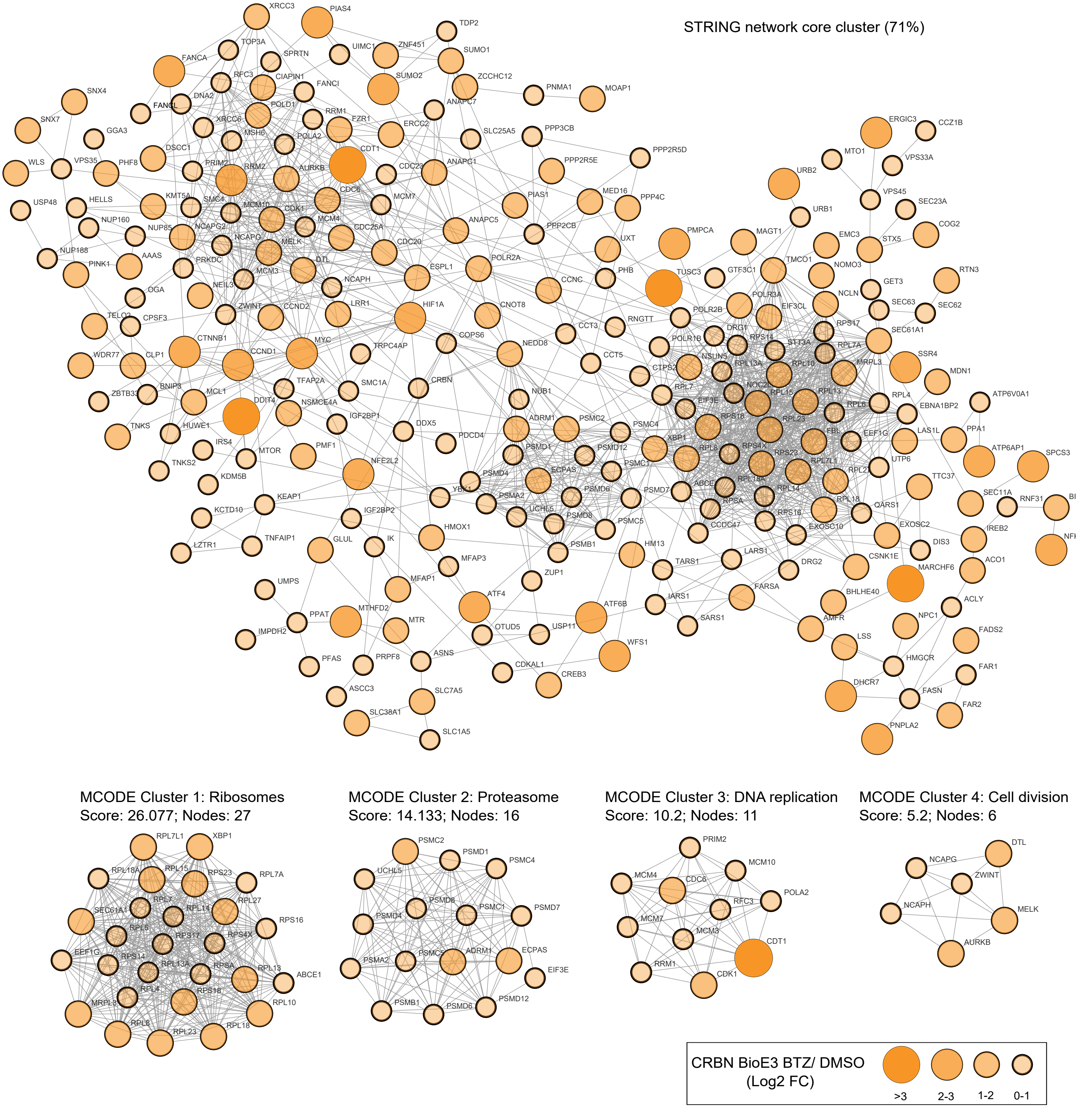

### Supplementary figure_Fig_S4

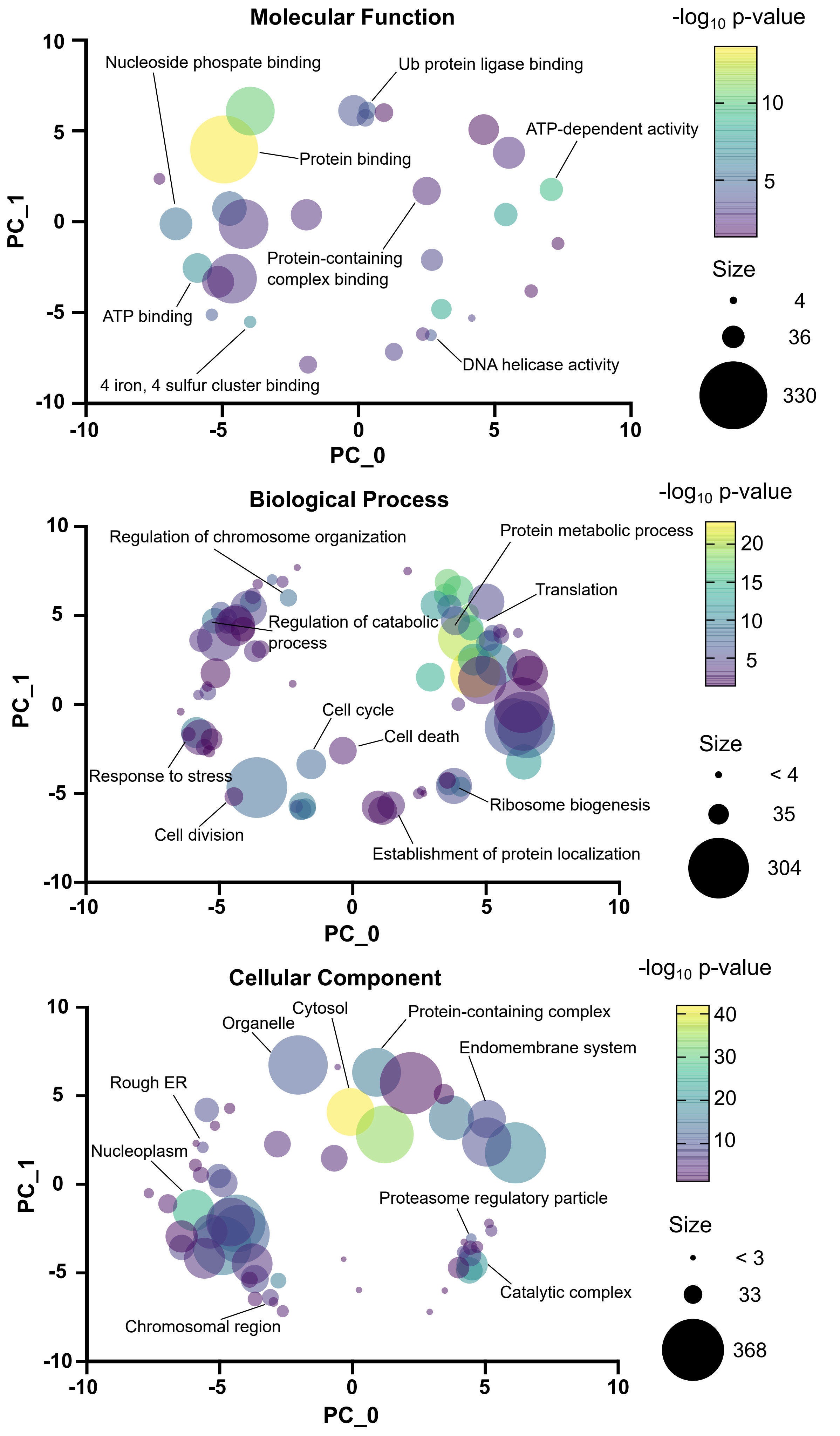


**CRBN BioE3 BTZ/ DMSO**

### Supplementary figure_Fig_S5

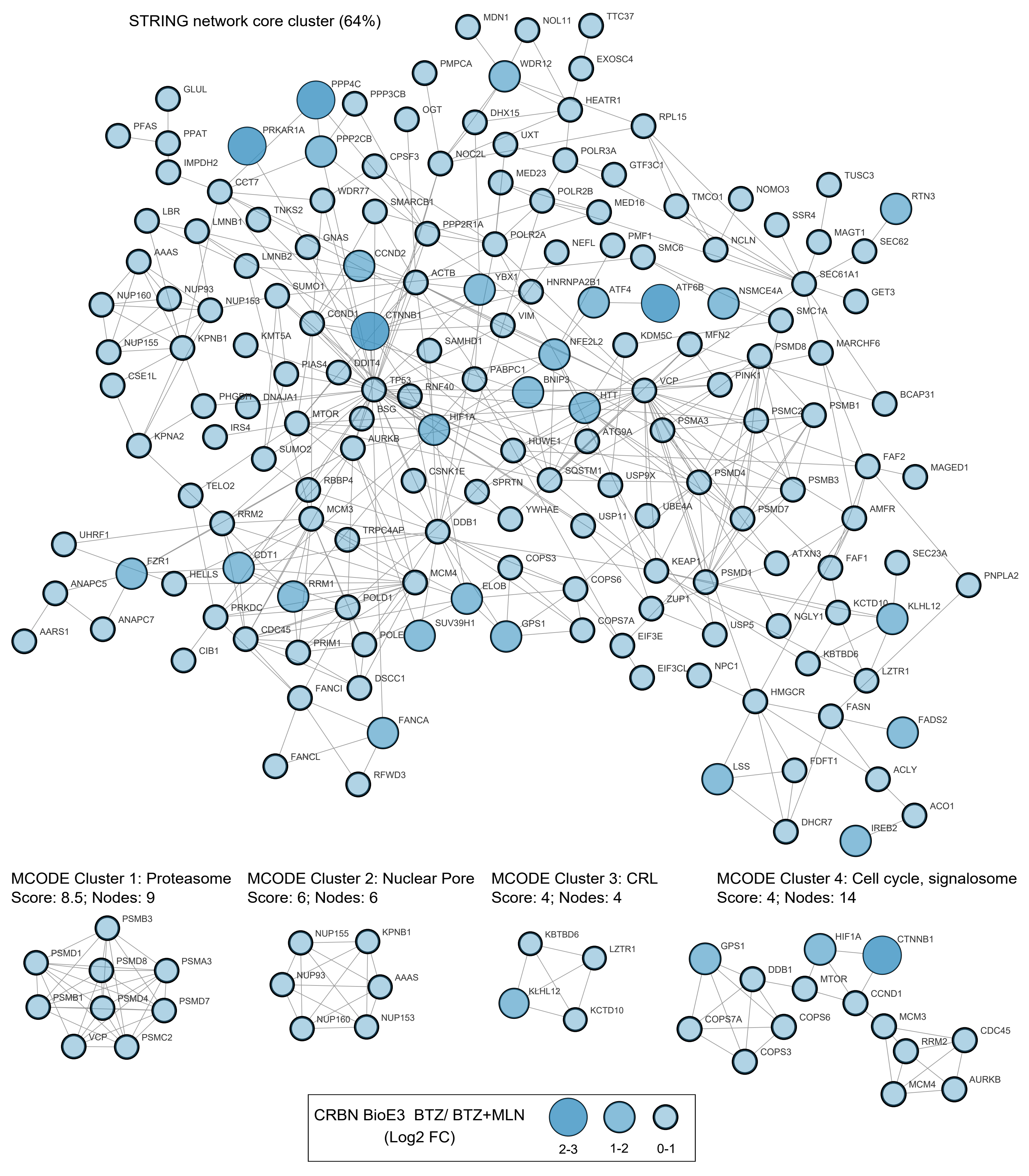

### Supplementary figure_Fig_S6

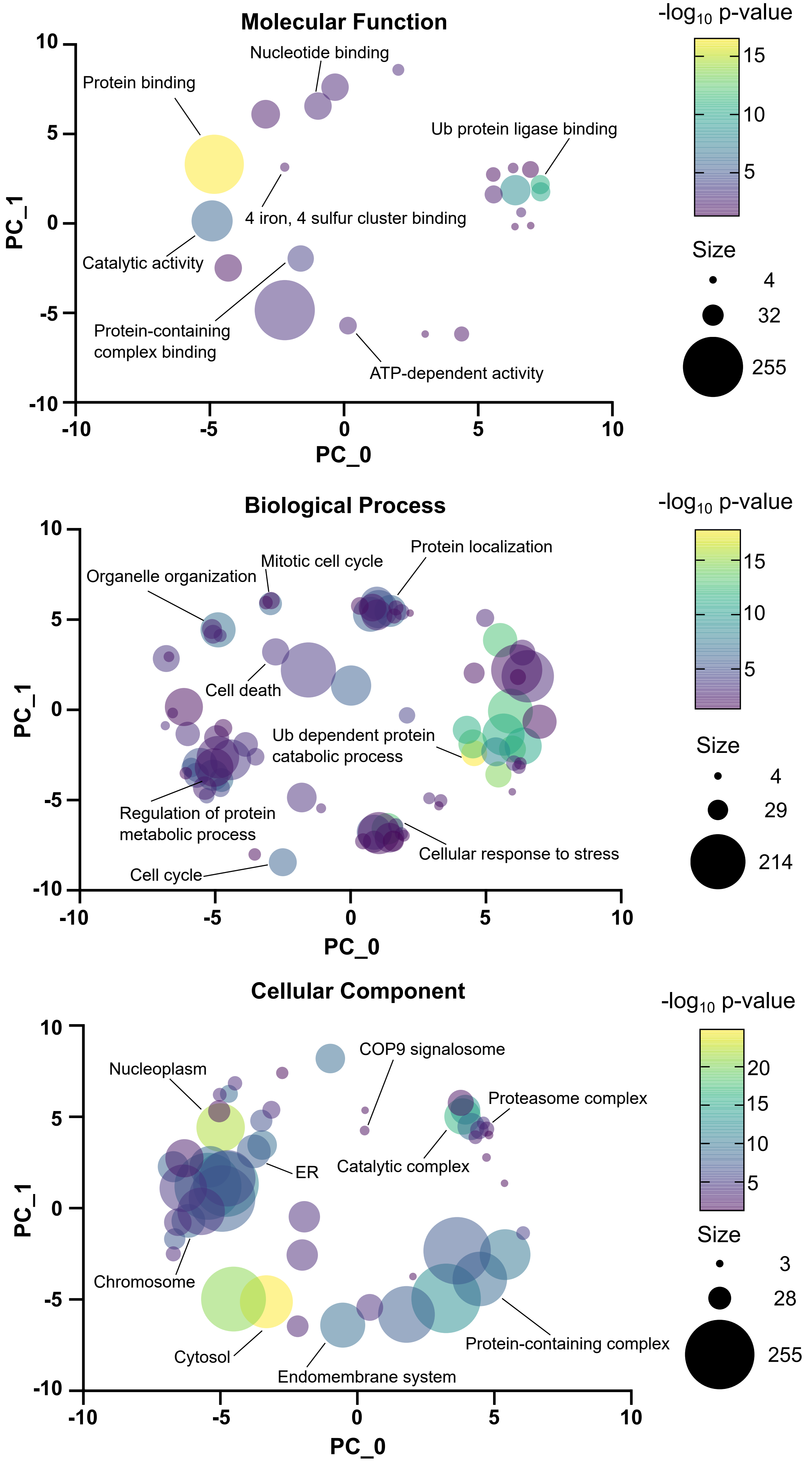


**CRBN BioE3 BTZ/ BTZ+MLN**

### Supplementary figure_Fig_S7

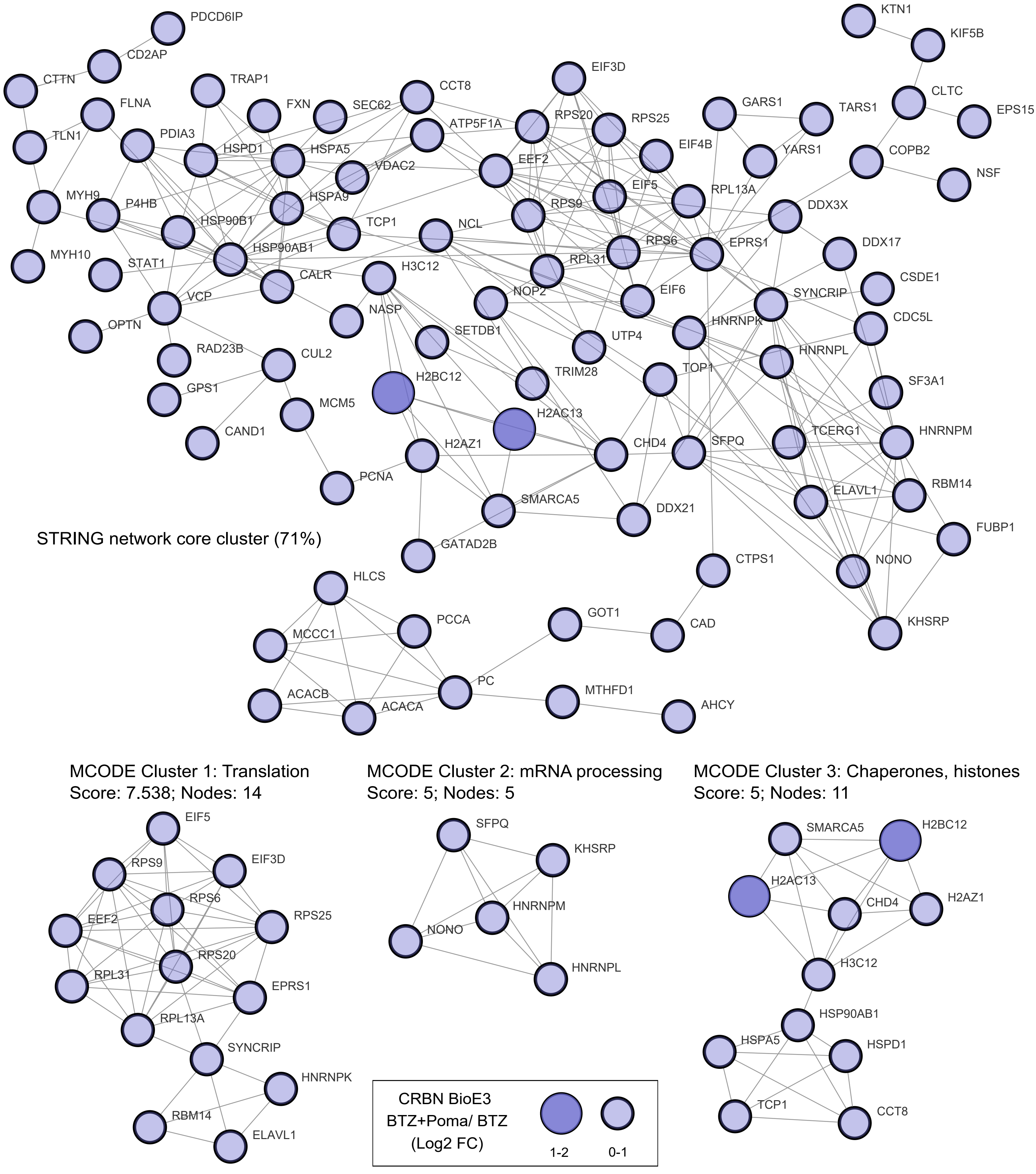

### Supplementary figure_Fig_S8

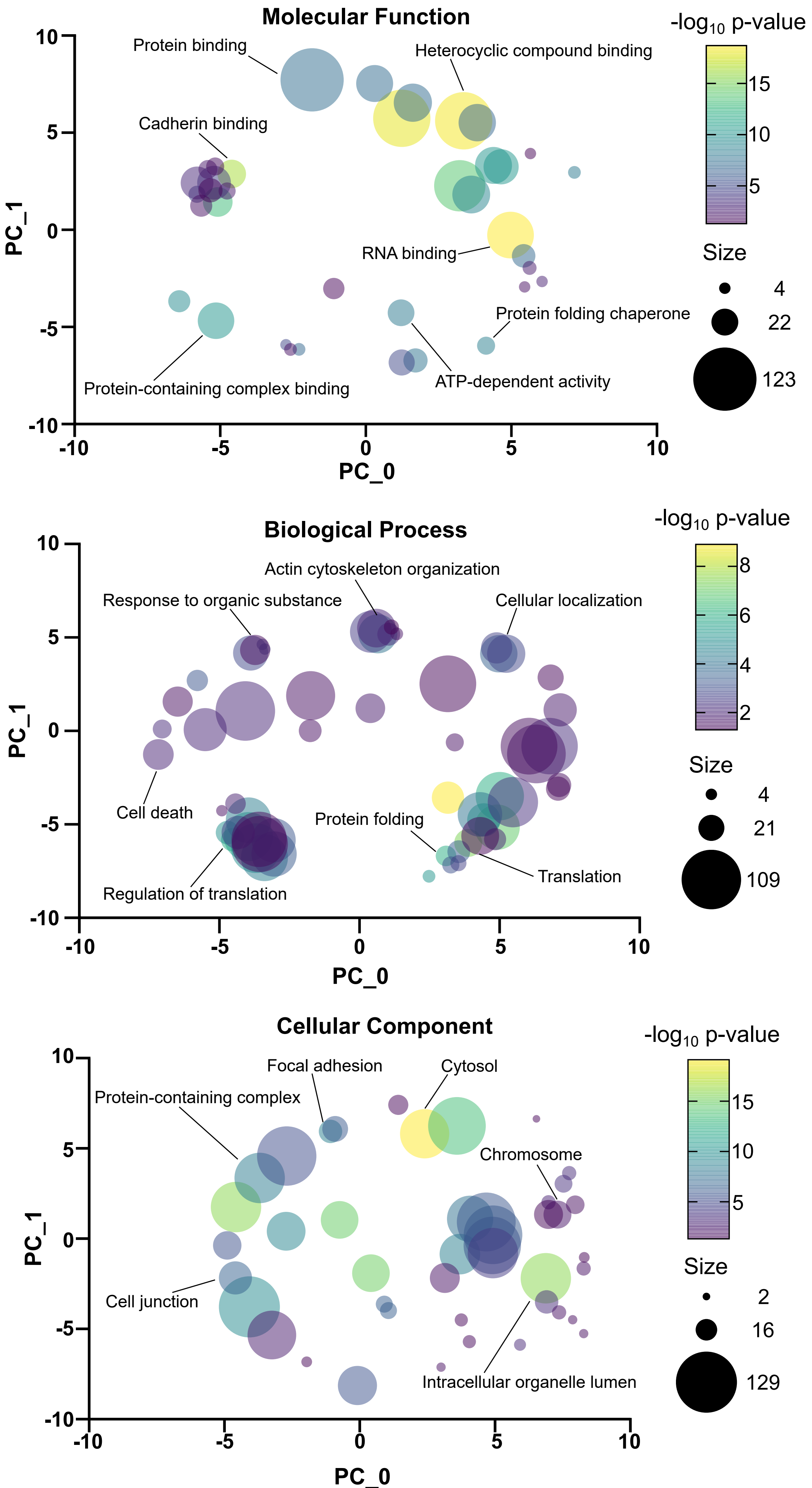


CRBN BioE3 Poma+BTZ/ BTZ
